# Host’s genetic background determines the outcome of reciprocal faecal transplantation on life-history traits and microbiome composition

**DOI:** 10.1101/2022.10.03.510653

**Authors:** Heli Juottonen, Neda N. Moghadam, Liam Murphy, Johanna Mappes, Juan A. Galarza

**Author notes:** equal contribution (Heli Juottonen, Neda N. Moghadam).

## Abstract

**Background:** Microbes play a role in fundamental ecological, chemical, and physiological processes of their host. Host life-history traits from defence to growth are therefore determined not only by abiotic environment and genotype but also by microbiota composition. However, the relative importance and interactive effects of these factors may vary between organisms. Such connections remain particularly elusive in Lepidoptera, which have been argued to lack a permanent microbiome and have microbiota primarily determined by their diet and environment.

We tested the microbiome specificity and its influence on life-history traits of two colour genotypes of the wood tiger moth (*Arctia plantaginis*) that differ in several traits, including growth. All individuals were grown in the laboratory for several generations with standardised conditions. We analysed the bacterial community of the genotypes before and after a reciprocal frass (i.e., larval faeces) transplantation and followed growth rate, pupal mass, and the production of defensive secretion.

**Results:** After transplantation, the fast-growing genotype grew at a significantly slower rate compared to the controls, but the slow-growing genotype did not change its growth rate. The frass transplant also increased the volume of defensive secretions in the fast-growing genotype but did not affect pupal mass. Overall, the fast-growing genotype appeared more susceptible to the transplantation than the slow-growing genotype.

Microbiome differences between the genotypes strongly suggest genotype-based selective filtering of bacteria from the diet and environment. A novel cluster of insect-associated *Erysipelotrichaceae* was exclusive to the fast-growing genotype, and specific *Enterococcaceae* were characteristic to the slow-growing genotype. These *Enterococcaceae* became more prevalent in the fast-growing genotype after the transplant, which suggests that the slower growth rate was potentially related to their presence.

**Conclusions:** We show that some genotype-specific life-history traits in a lepidopteran host can be reversed by a reciprocal frass transplantation. The results indicate that genotype-specific selective filtering can fine-tune the bacterial community at specific life stages, particularly the larval gut, even against a background of a highly variable community with stochastic assembly. Altogether, our findings suggest that the genotype of the host can influence its susceptibility to be colonized by microbiota with impact on key life-history traits.

## Background

Variation of traits within a population can be partly determined by genetic polymorphisms. Uncovering genotype-phenotype associations allows the analysis of the evolution and adaptive advantages of the traits. It is increasingly recognized that phenotype may also be influenced by the microbiome, which all animals including insects possess (Engel & Moran 2013). In general, the microbiome can potentially influence the host’s life history and fitness (Spor et al. 2011, Macke et al. 2017, Gould et al. 2018). In insects, the microbiome has been related to behavioural, nutritional, and life-history traits (Chaston et al. 2016, Vernier et al. 2020, Walters et al. 2020). Moreover, microbiome composition can vary according to the host’s genetic background (Macke et al. 2017, Walters et al. 2020). For instance, the gut microbiome can mediate the genotype effects on the phenotype: In *Drosophila*, the host genotype influences the microbiome composition, leading to differences in nutrition between phenotypes (Chaston et al. 2016).

A stable, symbiotic microbiome can confer benefits on the insect host, such as aiding its growth (Kikuchi et al. 2007, Douglas 2015, Hosokawa et al. 2016). On the other hand, transient, pathogenic bacteria could disadvantage the host (Holt et al. 2015, Smee et al. 2017), and even mutualistic symbionts can incur costs (Oliver et al. 2006). The outcome of these associations can depend on genotype-genotype interactions between the microbe and its host (Parker et al. 2017) as well as among microbes (Smee et al. 2021, Li et al. 2022). For example, in the pea aphid (*Acyrthosiphon pisum*), the host’s genotype influences the protection given by bacterial symbionts against pathogens (Parker et al. 2017, Weldon et al. 2020). In turn, host-to-microbe effects can play an important role in microbiome assembly within the host (Foster et al. 2017). Selective mechanisms that impact the establishment of microbes in insects include specialized organs (Ohbayashi et al. 2015, Lanan et al. 2016) and mechanisms that vary with host genetic background such as innate immunity (Lazzaro et al. 2006, Nyholm & Graaf 2012). Such filtering due to host traits and genetic background could influence the host’s fitness and life-histories (Koskella & Bergelson 2020).

The gut bacteria of lepidopteran larvae show metabolic potential to benefit the host by digesting and detoxifying food plants (Visôtto et al. 2009b, Xia et al. 2020) and by producing antimicrobial compounds against invaders (Shao et al. 2017). However, disruptions of the gut during moulting and metamorphosis, a highly alkaline pH (up to 11-12), lack of specialized gut structures, and fast passage of food can constrain the development of a consistent symbiotic microbiome (Engel & Moran 2013). Accordingly, several studies have concluded that there is no stable microbiome in Lepidoptera (Hammer et al. 2017, Staudacher et al. 2016, Martínez-Solís et al. 2020). Despite reports on the effects of diet, habitat, and developmental stage on gut bacteria (Broderick et al. 2004, Shao et al. 2014, Staudacher et al. 2016, Shao et al. 2017, Jones et al. 2019, González-Serrano et al. 2020), no clear consensus exists on the ecological roles of bacteria in Lepidoptera (Panigua Voirol et al. 2018).

Lepidopteran larval growth has been found to be correlated (Ruokolainen et al. 2016) and not correlated (Chatuverdi et al. 2017) with microbiome composition. Antibiotic treatment of lepidopteran larvae has similarly led to either increased (Visôtto et al. 2009a, Galarza et al. 2021) or decreased growth (Xia et al. 2020). Thus, the causal connections of microbiota and Lepidoptera growth traits remain elusive. One way to identify such connections is through microbiota transplants. Transplants (or bacteriotherapy) have been extensively applied in the biomedical field to study the potential of microbes in impacting health and disease (Zhang et al. 2018, Walter et al. 2018, Allegretti et al. 2019). The principle is to transfer microbes from a healthy individual to an unhealthy individual aiming to enrich beneficial microbes and restore a balanced microbiome. In insects, gut microbiota transplants are starting to reveal the importance of microbes in host development, immune response, and survival in dung beetles, cockroaches, bumblebees and parasitoid wasps (Näpflin and Schmid-Hempel 2016, Pietri et al. 2018, Jahnes et al. 2019, van Opstal and Bordenstein 2019, Parker et al. 2021). However, such studies are lacking in Lepidoptera, one of the most speciose and ecologically important groups of insects, in which less than 0.1% of species have been screened for microbes (Panigua Voirol et al. 2018). The uncertainties of the functional role and specificity of microbes in Lepidoptera make this group a particularly important target for transplant experiments.

Here, we investigate the effect of microbiome transplantation on wood tiger moths (*Arctia plantaginis*) with distinct genetic background and life-history traits. A reciprocal faecal transplantation was carried out between wood tiger moths of two colour genotypes that differ in the duration of their larval stage by adding frass (i.e., larval faeces) to their diet. The experimental insects have been reared in the laboratory for several generations, kept in similar conditions and fed the same diet. We followed the microbiome compositional changes during larval development and in the resulting adults. Thus, any consistent microbiome differences between host genotypes could reflect the selective filtering of bacteria. We ask i) if each genotype has its own associated microbiome, ii) if it is stable across life-stages, and iii) if the microbiome transplant can reverse growth rate between the genotypes. We also examine if the transplant impacts other important fitness traits such as pupal mass and the volume of defensive secretions.

## Methods

### Study species and genotype lines

The wood tiger moth is an aposematic species distributed throughout the Holarctic (Hegna et al. 2015). In Europe, males display colour polymorphism, having yellow or white hindwings and co-occur at variable frequencies within populations (Galarza et al. 2014, Hegna et al., 2015). The two colour morphs differ in key fitness traits such as mating success (Nokelainen et al. 2012, Gordon et al. 2015), immune responses (Nokelainen et al. 2013), protection against predators (Lindstedt et al. 2011, Rojas et al. 2019), and flight activity (Rojas et al. 2015). The yellow-white hindwing polymorphism is determined by a single Mendelian locus with two alleles in which the yellow allele (y) is recessive to the white (W) allele (Nokelainen et al. 2022). Hence, the white colouration is produced by WW and Wy allelic combinations, whereas yy produces yellow. Analyses of selection lines show that the homozygous genotypes differ in the length of their larval stage (i.e., from egg hatching to pupation).

Individuals of the WW genotype have a significantly shorter larval stage than those of the yy genotype (Fig. S1). Adults of both colour morphs release defensive secretions from their anal cavity which are effective at deterring invertebrate predators, yellows having stronger chemical defence than whites (Rojas et al. 2017). Thus, this species offers a good opportunity to study the impact of microbes on life-history and fitness traits in relation to the host’s genetic background.

### Larval sampling and rearing before frass transplant

Genotype selection lines of wood tiger moths have been maintained for over 12 generations at the University of Jyväskylä, Central Finland. For this study we selected four families of WW and four of yy genotype with a mean of 239 eggs/family to characterize their bacterial communities as they develop with or without faecal transplantation (see below). We included four families to cover for possible variation among families in the analysis of genotype effects, and we did not analyse family effects. The general rearing protocol and the pedigree are described in detail in Nokelainen et al. (2022) and DePasqual et al. (2022). Here we modified the rearing protocol as follows. Immediately after hatching and before being given any food, we collected larvae to assess the bacteria in newly hatched larvae. Newly hatched larvae are too small for dissection (∼2 mm), and hence, the whole larva was used. The larvae were surface sterilized to exclude microbial contamination from the environment. We cut the filter from a 1-ml filter tip and placed it inside a 1.5-ml tube. We pooled four larvae into a sample on the filter and added 450 μl of autoclaved double-distilled water (AddH_2_O) creating a whirlpool with a 1-ml filtered pipette tip for 2 minutes. The water was collected and the procedure was repeated three times. The collected **washing water** (Table 1) was stored at −20 °C until DNA extraction to represent bacteria on the outside of the larva including environmental contamination. The washed larvae were then transferred to a new 1-ml filter tip and rinsed with 450 μl of 5% sodium hypochlorite (NaOCl) solution and centrifuged at 2000 rpm for 30 seconds. The rinsing process was repeated three times, after which the surface sterilized larvae were stored at −20 °C until DNA extraction (**hatched larvae** in Table 1).

**Table 1.**
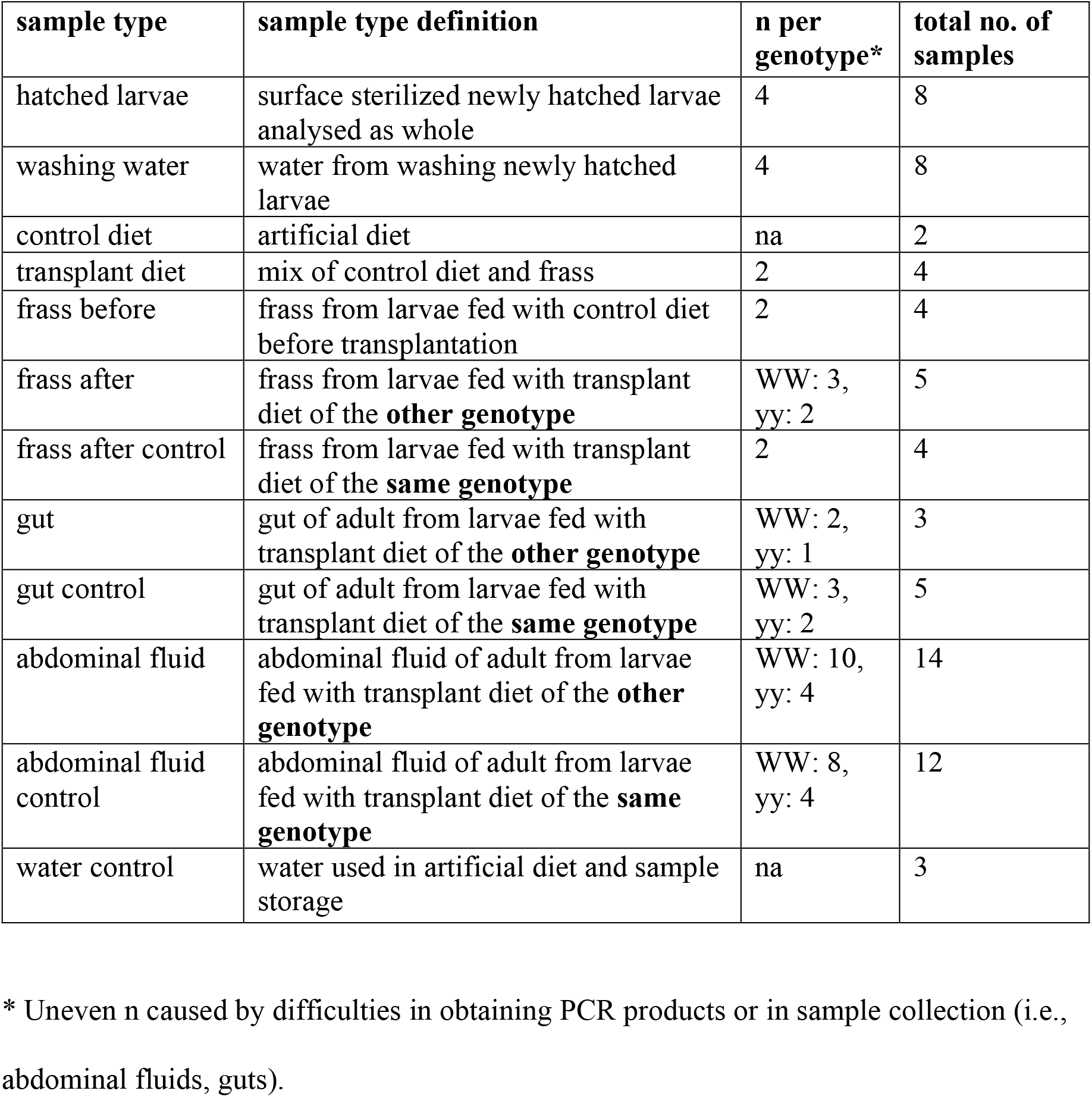
Sample types included in bacterial 16S rRNA gene sequencing.

| sample type | sample type definition | n per genotype* | total no. of samples |
| --- | --- | --- | --- |
| hatched larvae | surface sterilized newly hatched larvae analysed as whole | 4 | 8 |
| washing water | water from washing newly hatched larvae | 4 | 8 |
| control diet | artificial diet | na | 2 |
| transplant diet | mix of control diet and frass | 2 | 4 |
| frass before | frass from larvae fed with control diet before transplantation | 2 | 4 |
| frass after | frass from larvae fed with transplant diet of the **other genotype** | WW: 3, yy: 2 | 5 |
| frass after control | frass from larvae fed with transplant diet of the **same genotype** | 2 | 4 |
| gut | gut of adult from larvae fed with transplant diet of the **other genotype** | WW: 2, yy: 1 | 3 |
| gut control | gut of adult from larvae fed with transplant diet of the **same genotype** | WW: 3, yy: 2 | 5 |
| abdominal fluid | abdominal fluid of adult from larvae fed with transplant diet of the **other genotype** | WW: 10, yy: 4 | 14 |
| abdominal fluid control | abdominal fluid of adult from larvae fed with transplant diet of the **same genotype** | WW: 8, yy: 4 | 12 |
| water control | water used in artificial diet and sample storage | na | 3 |

\* Uneven n caused by difficulties in obtaining PCR products or in sample collection (i.e., abdominal fluids, guts).

The remaining larvae (374/genotype) were split into groups of 20-25 larvae and reared inside sterile petri dishes. The dishes were kept in climate chambers in a 18:6 h light:dark cycle at 21 °C during the light and 14 °C during the dark. An artificial diet was prepared consisting of 3 g agar, 32.1 g semolina, 8.58 g yeast, 8.3 g wheat germ, 1.76 g Vanderzant vitamin mix, 1.8 ml nipagin and 180 μl acetic acid in 200 ml freshly boiled AddH_2_O to minimize diet-derived bacteria. Two samples of this diet were stored at −20 °C until DNA extraction (**control diet** in Table 1). Roughly 5 g of this diet was presented to the larvae on top of a sterilized microscope slide inside the petri dish. After 48 h, approximately 0.5 g of larval faeces, hereafter frass, was collected from the bottom of the petri dish using sterilized tweezers and stored at −20 °C until DNA extraction (**frass before** in Table 1).

### Frass transplantation and rearing

To prepare diets for frass transplantation between genotypes, approximately 10 frass pellets from each petri dish were collected after 48 h of giving the **control diet** and pooled to obtain ∼1 g/genotype and mixed with 50 g of the control diet. Four samples of this transplant diet (two of each genotype) were mixed with 2 ml of boiling AddH_2_O and stored at −20 °C until DNA extraction (**transplant diet** of genotypes WW and yy in Table 1).

The larvae, all in their 3^rd^ or 4^th^ instar, were then divided into treatment and control groups in sterile petri dishes with 10-15 larvae of the same genotype per petri dish. The treatment group was fed the opposite genotype’s transplant diet: each petri dish of WW larvae received ∼5 g of yy transplant diet, and each petri dish of yy larvae received ∼5 g of WW transplant diet. The control group larvae received ∼5 g of transplant diet of their own genotype. The petri dishes were kept in the climate cabinets in the same conditions as above. Twenty-four hours after the food was given, approximately 1 g of frass was collected from each genotype as above and stored at −20 °C until DNA extraction (**frass after, frass after control** in Table 1). The rearing continued until all larvae pupated or died. The pupae were placed individually in 150-ml plastic containers, kept in the climate chambers in the same conditions and weighed to the nearest milligram. From a subset of the emerging adults, we dissected the gut following Moghadam et al. (2018) with minor modifications. Briefly, each adult moth was placed on a sterile petri dish and its head was removed using a sterilized scalpel. A drop of AddH_2_O was placed next to the abdomen and the gastrointestinal tract (i.e., gut) including the crop, foregut, midgut, and hind gut was pulled out using sterilized forceps under a light stereoscope with a Bunsen burner next to it to reduce the risk of contamination. The dissected guts were placed individually in 30 μl of AddH_2_O and stored at −20 °C until DNA extraction (**gut** in Table 1). Likewise, we collected the abdominal defensive secretions from the adults. We gently pressed the abdomen of live adults with sterilized tweezers until the secretion was released from the anal cavity. The secretion was collected using UV-sterilized 10-μl glass capillaries under a laminar flow, measured with a digital caliper, and placed individually in 30 μl of AddH_2_O and stored at −20 °C until DNA extraction (**abdominal fluid** in Table 1). Finally, we took 30 μl of the AddH_2_O batch used to prepare all the samples above and stored it at −20 °C until DNA extraction (**water control** in Table 1).

### Life histories

We followed several life-history traits of individual larvae, pupae, and adults from the different genotypes and treatments. The overall developmental rate was determined by counting the number of days elapsed from egg hatching until adult eclosion. This included the larval and pupal stages. We further analysed the development rate within the larval stage (i.e., from egg hatching until pupation), as well as within the pupal stage (i.e., from pupation to adult eclosion). In addition, at the pupal stage, we recorded the weight of all individual pupa, and at the adult stage, we measured the volume of abdominal defensive secretions as described above.

### DNA extraction, PCR and PacBio amplicon sequencing

DNA was extracted by homogenizing the sample (larvae, frass, gut, abdominal fluid, diet) in 30 μl of water with a metal bead (Ø 2.3mm) in a Bead Ruptor (OMNI) at speed 3.93 m/s for 2 × 30 s. After homogenization, the samples were boiled at 100 °C for 10 min and stored at −20 °C until further use. DNA quantification was performed with the Qubit BR DNA kit (ThermoFisher).

To assess the bacterial diversity in the larvae, their frass, and adult moths, we amplified ∼1550 bp of the 16S ribosomal RNA (rRNA) gene using custom primers (forward 5’-AGAGTTTGATCMTGGCTCAG-3’, reverse 5’-CCTTGTTACGACTTCACCCCAG-3’). The primers were designed using Primer3 (Untergasser et al. 2012) from Lepidoptera-associated 16S rRNA gene sequences downloaded from National Center for Biotechnology Information (NCBI) and aligned using ClustalW (Sievers et al. 2011). Polymerase chain reactions (PCR) were performed in a C1000 thermal cycler (Bio-Rad) using 5 μl of DNA template, 1 × Platinum SuperFI Mastermix (ThermoFisher), 1 × enhancer buffer, 0.5 μM of each primer, and 0.5 mM MgCl_2_ in reaction volume of 20 μl. The cycling conditions were as follows: 98 °C for 30 s, 40 cycles of 98 °C for 10 s, 49 °C for 10 s, 72 °C for 1 min, and a final elongation of 5 min at 72 °C. The PCR products were run in 3% agarose gels and the bands were excised using gel cutting tips (Axygen) and purified by centrifuging through 1-ml filter tips at 6000 rpm for 15 min. The purified PCR products’ concentration was measured using PicoGreen dsDNA Assay Kit (ThermoFisher), and a sequencing library was prepared according to the PacBio multiplexed amplicon library preparation protocol. The samples were barcoded and sequenced in a PacBio Sequel at the Novogene sequencing laboratories. In addition, a mock community (ZymoBIOMICS Microbial Community DNA Standard, Zymo Research) and a water control (i.e., negative control) were amplified and sequenced with each library.

### Sequence processing and quality control

Sequence reads were processed with PacBio tools distributed in Bioconda. PacBio subreads were combined into consensus sequences with ccs in package pbccs (v.6.0.0) with the default settings. The consensus sequences (797984 reads) were demultiplexed based on barcodes with lima (v. 2.0.0) with the settings --peak-guess, --different- -ccs, --min-length 1440, -- max-input-length 1580, --min-end-score 26, and --split-bam-named. The resulting bam files were converted into fastq with bam2fastq in the package bam2fastx (v. 1.3.1). Sequences were submitted to the National Center for Biotechnology Information Sequence Read Archive under accession code PRJNA804133.

The reads were processed further and amplicon sequence variants (ASVs) inferred in DADA2 (v. 1.16.0, Callahan et al. 2016) following guidelines for PacBio data (Callahan et al. 2019, https://benjjneb.github.io/LRASManuscript/LRASms_fecal.html) in R (v. 4.0.4, R Core Team 2021) and RStudio (v. 1.4.1106). Primers were removed with the command removePrimers. Reads were filtered with the command filterAndTrim and the settings minQ=3, minLen=1300, maxLen=1600, maxN=0, and maxEE=2. The reads were dereplicated and denoised using PacBio-specific error estimation function. Chimeras were removed from the denoised reads with the command removeBimeraDenovo and setting minFoldParentOverAbundance=3.5. Taxonomy was assigned against the Silva database (v. 138.1, Quast et al. 2013). The ASV data was imported into phyloseq (v. 1.32.0, McMurdie and Holmes, 2013), and ASVs that were assigned into chloroplasts or mitochondria or not assigned to Bacteria were removed. Potential contaminants were examined based on ASVs in water control and washing water samples (Table 1) with the package decontam based on frequency (threshold 0.2), prevalence (threshold 0.5), and inspection of frequency vs. DNA concentration plots (Davis et al. 2018), and none were detected. This resulted in 226 ASVs (when excluding ASVs in the mock community controls) and on average 6898 reads per sample (in total 565705 reads).

### Statistical analyses and phylogenetic diversity measures

All analyses were carried out in R (v. 4.0.4) through RStudio. The package ggplot2 (Wickham 2016) was used for generating plots. To examine the effect of the frass transplantation on the life histories of the genotypes, we implemented a Kruskal-Wallis one-way analysis of variance (ANOVA) followed by pairwise Dunn’s tests to the traits measured. All tests’ significance values were corrected for multiple comparisons. This non-parametric approach was chosen because the samples violate parametric assumptions of normality and/or equality of variance (Fig. S2).

For comparing bacterial diversity and community composition, the ASV table was rarefied to the median number of reads (6915 reads) with the function rrarefy in vegan (v. 2.5.7, Oksanen et al. 2020). If a sample had fewer reads than the median, all its reads were included. Then, the ASV table was standardized to relative abundances. Sequences of the ASVs were aligned and a phylogenetic tree was constructed using RAxML (model GRT+gamma, Stamakis 2014) on the SILVA ACT server (Pruesse et al. 2012). Faith’s phylogenetic diversity (PD) and ASV richness were determined in picante (v. 1.8.2, Kembel et al. 2010). The phylogenetic tree was converted into a phylogenetic distance matrix with the function cophenetic. Measures of phylogenetic relatedness among communities (phylogenetic beta diversity) were calculated in picante as mean pairwise distance (MPD, Webb et al. 2002, function comdist, abundance weighted) and mean nearest taxon distance (MNTD, function comdistnt, abundance weighted). MPD emphasizes the clustering of basal clades in the phylogenetic tree, whereas MNTD emphasizes patterns closer to the tips of the tree. These values were used as distance measures in non-metric multidimensional scaling (NMDS) with function metaMDS in vegan and permutational multivariate analysis of variance (PERMANOVA) with function adonis2 in vegan (Anderson 2001).

Separate phylogenetic trees of *Erysipelotrichaceae* and *Enterococcus* ASVs were constructed by aligning the sequences and selected reference sequences (described strains and similar environmental sequences identified by Blast searches) with SINA v. 1.2.11 on the SILVA ACT server and inferring a maximum likelihood tree with RAxML (model GTRGAMMA, Stamakis 2014) in QIIME2 (v. 2021.8.0, Bolyen et al. 2019).

We used the same phylogenetic relatedness measures as above to assess phylogenetic clustering of bacterial communities against null models across sample types. This analysis aims to determine whether bacterial community assembly and turnover across life stages are driven by niche-based (i.e., environmental filtering) or stochastic processes (Stegen et al. 2012). For example, if gut conditions favour the proliferation of specific bacterial taxa, it would be shown as phylogenetic clustering. Net relatedness index (NRI) was calculated as the standardized effect size of MPD (function ses.mpd in picante, abundance weighted) multiplied by −1. Nearest taxon index (NTI) was calculated as standardized effect size of MNTD (function ses.mntd, abundance weighted) multiplied by −1. Positive values of NRI and NTI indicate phylogenetic clustering (taxa are more closely related than by chance), whereas negative values indicate phylogenetic overdispersion (taxa are less closely related than by chance). Values of NRI and NTI differing from 0 and thus showing more clustering by chance were identified by Student’s t-test (function t.test). To compare mechanisms of phylogenetic turnover along life history and experimental stages, we calculated βNTI values for pairs of communities according to Stegen et al. (2012) and R scripts from https://github.com/stegen/Stegen_etal_ISME_2013. βNTI > 2 indicates significantly higher community turnover than by chance driven by deterministic selection (Stegen et al. 2012, 2013). βNTI between −2 and 2 indicates community assembly driven by stochastic processes. βNTI < −2 indicates less community turnover than by chance.

## Results

### Life-history effects of frass transplantation

The overall developmental time (i.e., from egg hatching to adult eclosion) differed between the controls of genotypes WW and yy (Fig. S3), as in the stock selection lines (Fig. S1). This difference was mainly due to faster growth of WW individuals at the larval stage (Fig. 1). The length of the pupal stage did not differ between the genotypes (Kruskal-Wallis one-way ANOVA statistic = 7.74, P > 0.05) (data not shown). When WW larvae received the frass transplant from genotype yy, they grew slower than the control WW larvae that received WW frass (Fig. 1, Fig. S3). The opposite was not observed: the WW frass transplant had no effect on the growth of yy larvae.

**Fig. 1.**
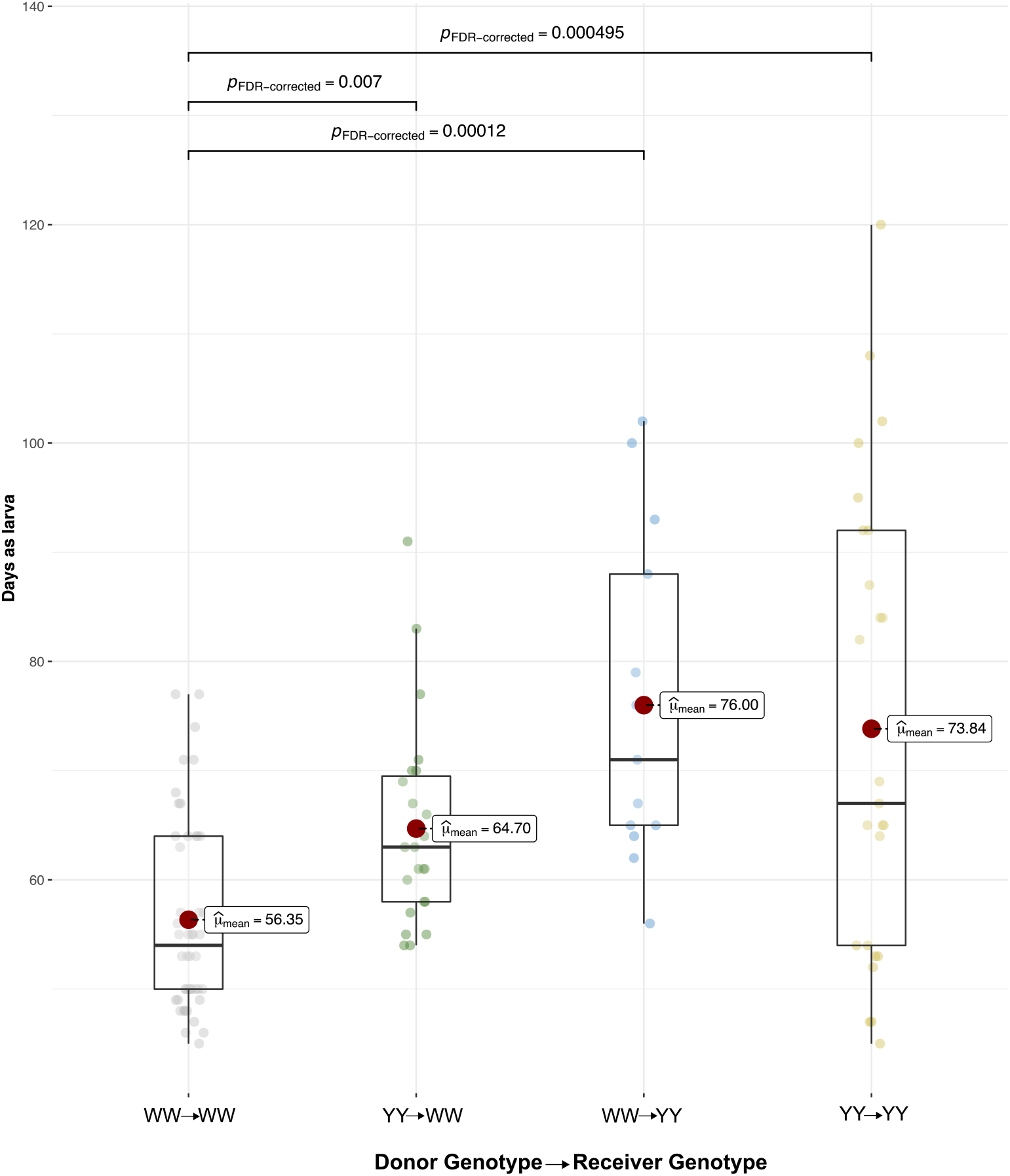
Days as larvae of *A. plantaginis* in reciprocal frass transplantation between genotypes WW and yy and in control transplantations within the genotypes.

Pupal weight differed with genotype in the controls, with WW pupae being lighter than yy pupae (Fig. S4). However, we observed no differences in pupal weight between the controls and the pupae that received the frass transplant of the other genotype. This suggests that the frass transplant did not affect pupal weight.

The adults of genotype WW secreted a smaller volume of defensive abdominal fluids than the adults of yy in the controls. Transplantation with yy frass significantly increased abdominal fluid secretion in WW adults (Fig. 2). As in the case of growth rate, this could point to greater susceptibility or adaptability of the WW genotype to the transplantation and the presence of foreign bacteria. Bigger volumes in defensive secretions were also observed in the yy genotype transplanted with WW frass. However, this difference was not significant relative to the yy controls, likely because of the unbalanced number of samples. Overall, it can be suggested that frass transplantation generally increased the volume of defensive secretions.

**Fig. 2.**
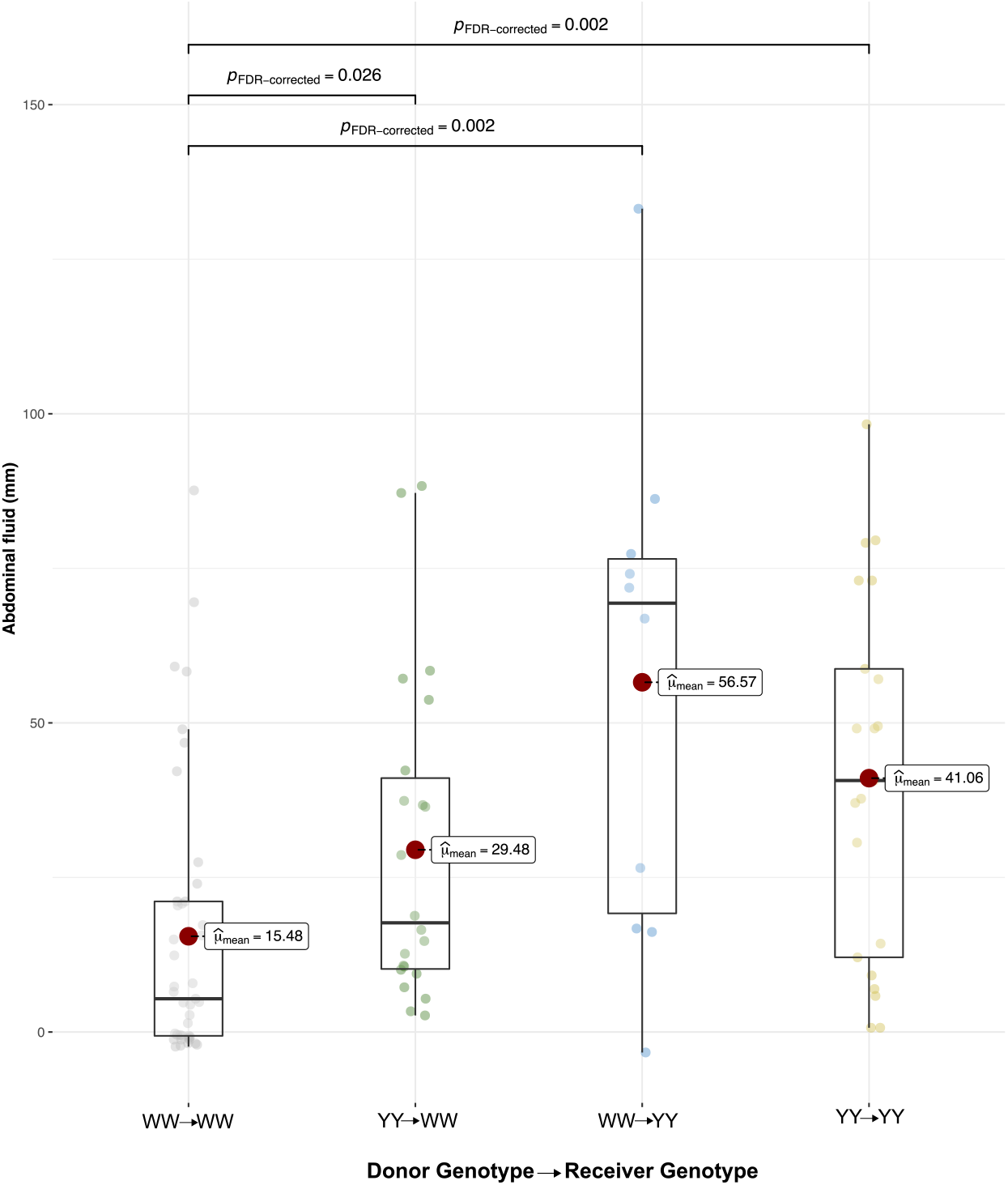
Secretion of defensive abdominal fluids of *A. plantaginis* adults in reciprocal frass transplantation between genotypes WW and yy and in control transplantations within the genotypes.

### Bacterial community composition and diversity with genotype, life stage and frass transplant

Bacteria detected in newly hatched larvae showed no difference in phylogenetic diversity or ASV richness between the genotypes (Fig. S5). Compared to newly hatched larvae, larval frass before transplantation had lower phylogenetic diversity (Fig. S5), which indicated that frass contained a reduced set of bacteria. However, the frass bacteria were not solely a subset of the bacteria detected in newly hatched larvae. Frass before transplantation shared only one out of five ASVs (genotype WW) or three out of 11 ASVs (genotype yy) with the newly hatched larvae (Fig. 3).

**Fig. 3.**
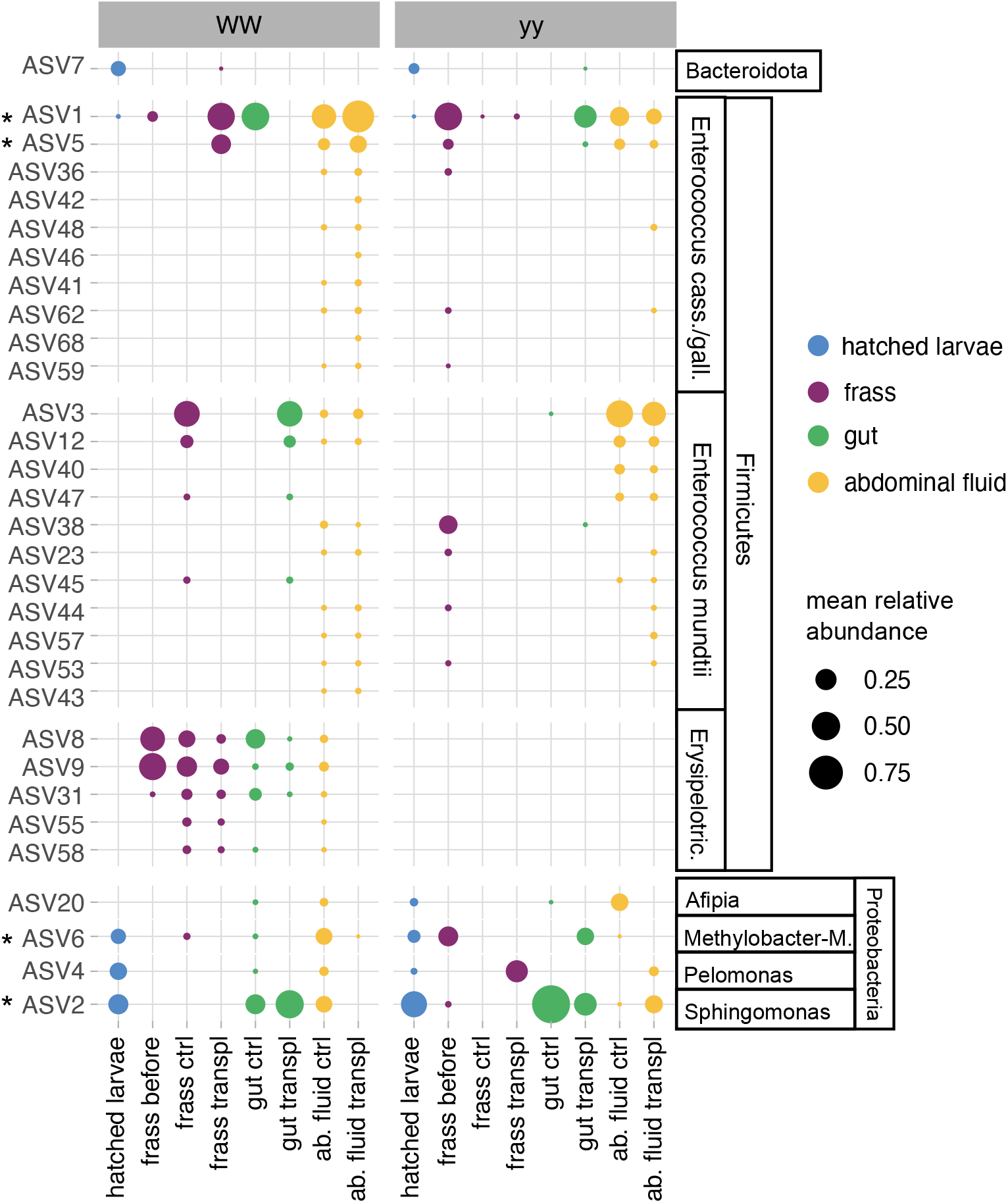
Distribution of bacterial 16S rRNA gene amplicon sequence variants (ASVs) in *A. plantaginis* genotypes WW and yy with life stage in frass transplantations between genotypes (transpl) and in control transplantation within genotypes (ctrl). Only ASVs occurring in more than two samples are shown. * marks ASVs with > 50% prevalence across the samples. Replicate samples within the same sample type and genotype have been merged and the mean relative abundance of ASVs is shown. Cass. stands for *casseliflavus*, gall. for *gallinarum*, Erysipelotric. for *Erysipelotrichaceae*, Methylobacter-M. for *Methylobacter-Methylorubrum*, and ab. fluid for abdominal fluid.

Unlike in newly hatched larvae, the bacterial community in frass differed between the genotypes (Fig. 4, Fig. S6; PERMANOVA hatched larvae R^2^ = 0.27, P = 0.24; frass R^2^ = 0.38, P = 0.001). Frass of genotype WW collected before transplantation had lower ASV richness and phylogenetic diversity than yy frass (Fig. S5). Bacteria in WW frass were dominated by *Erysipelotrichaceae* (*Firmicutes*; Fig. 5), which were only detected in the genotype WW: in addition to larval frass in WW adult gut and abdominal fluids (Fig. 3).

**Fig. 4.**
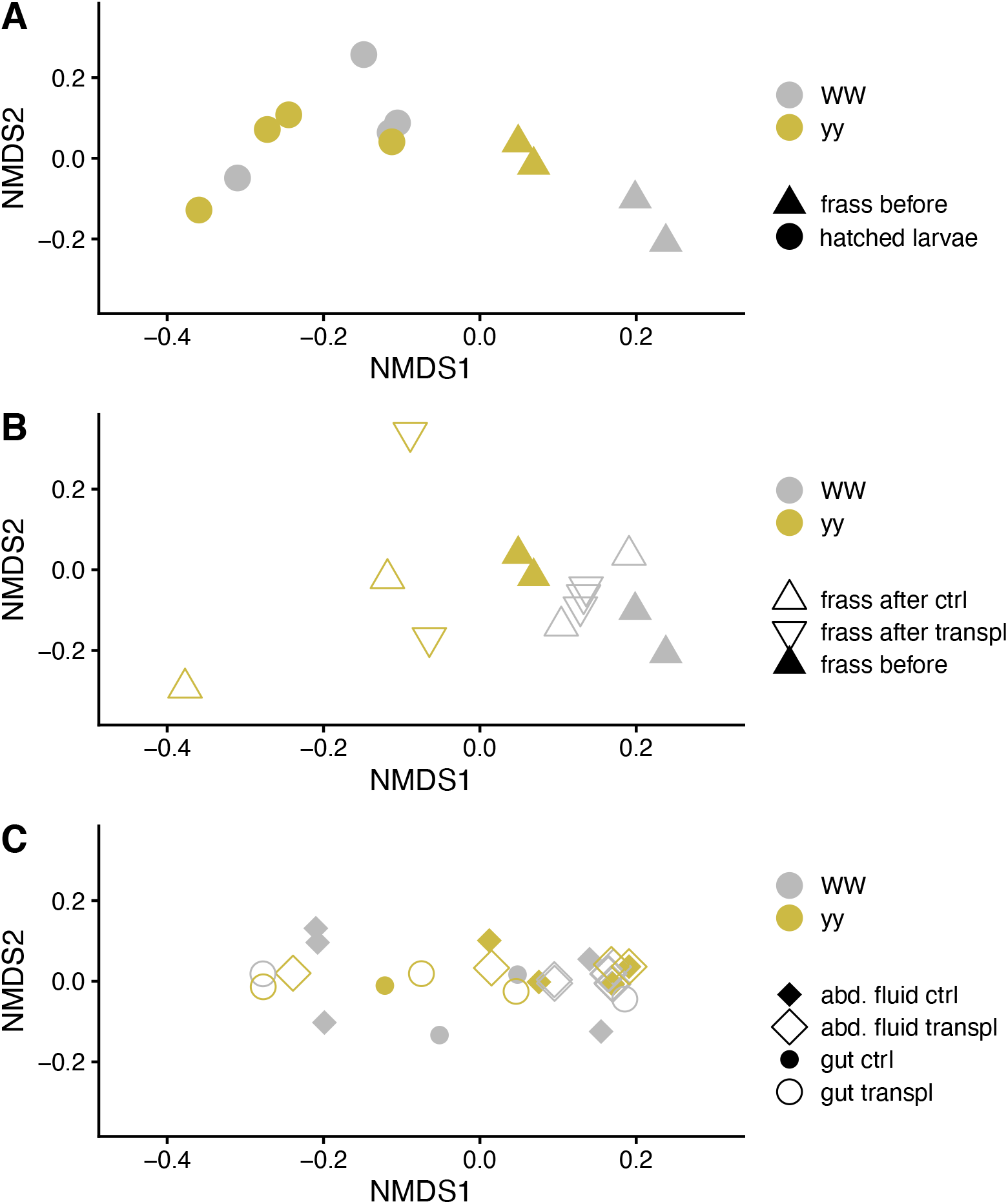
Non-metric multidimensional scaling (NMDS) plots of bacterial community in *A. plantaginis* genotypes WW and yy based on phylogenetic distances (MNTD, mean nearest taxon distance). A) Newly hatched larvae and frass before transplantation, B) frass before and after transplantation in controls within genotypes (ctrl) and between genotypes WW and yy (transpl), C) gut and abdominal fluid (abd. fluid). Panels A, B and C are based on the same ordination but plotted separately. Stress = 0.15.

**Fig. 5.**
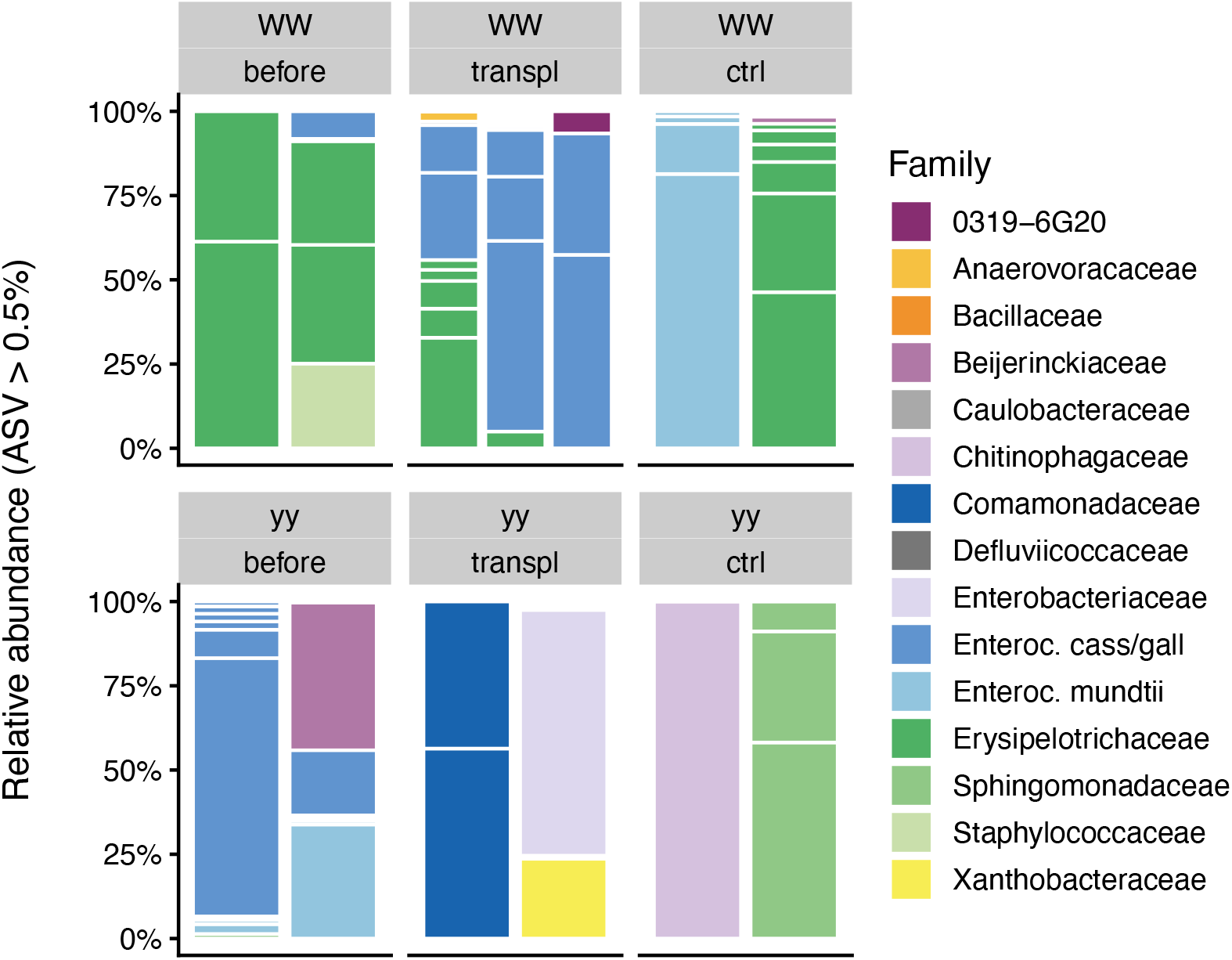
Taxonomic distribution of bacterial 16S rRNA gene amplicon sequence variants (ASVs) at family level in frass before and after transplantation within genotypes WW and yy (ctrl) and between genotypes WW and yy (transpl). White horizontal lines separate ASVs and each ASV with relative abundance above 0.5% is shown as its own section in the columns. Family ‘0319-6G20’ belongs to phylum *Bdellovibrionota*, class *Oligoflexia*. Enteroc. stands for *Enterococcus*, and cass/gall for *casseliflavus*/*gallinarum*.

These *Erysipelotrichaceae* belong to a novel cluster with no described species that includes uncultured bacteria detected in termite gut (Fig. S7; Hongoh et al. 2005, Mikaelyan et al. 2015). Frass of genotype yy was instead dominated by *Enterococcaceae* (*Firmicutes*; Fig. 5). These ASVs included ASV1 and ASV5 (Fig. 3) that occurred across the genotypes in frass, adult gut, and abdominal fluids and had 100% sequence similarity to *Enterococcus gallinarum* and *E. casseliflavus* strains from Lepidoptera (Fig. S8; Dantur et al. 2015, Chung et al. 2018). These *Enterococcaceae* ASVs were among the only four common core ASVs detected in one or more sample types with 50% prevalence (Fig. 3).

When WW larvae received the yy frass transplant, their frass retained the WW-specific *Erysipelotrichaceae*. Notably, WW frass also gained or showed an increased relative abundance of the two common *Enterococcaceae* ASVs detected in yy frass (ASV1, ASV5) (Fig. 5). The prominence of these ASVs in WW frass after transplantation suggests they may have originated from the yy frass transplant. *Enterococcus* ASVs became more prevalent also in one of the WW controls that received WW frass, but these ASVs represented a different cluster within *Enterococcus* not detected in yy frass (ASV3, ASV12, 99-100% sequence similarity to *E. mundtii* strains, Fig. S9). Bacteria in yy frass, on the other hand, changed almost completely both with WW frass transplant and in the controls that received genotype yy’s own frass (Fig. 4B, Fig. 5). After transplant, yy frass bacteria consisted of sporadic ASVs representing bacterial groups detected in the control diet and transplant diet fed to the larvae (Fig. 5, Fig. S9). Therefore, we found no evidence pointing to the transfer of WW frass bacteria to genotype yy. Variation of bacterial community in adult gut and abdominal fluid could not be linked to the transplant treatment or to genotype, partly due to large variation among individuals (Fig. 4C, Fig. S10). The ASVs affiliated with *E. casseliflavus/gallinarum* and *E. mundtii* occurred in adult gut with no consistent pattern. In adult abdominal fluids, both *Enterococcus* types were present in both genotypes, but *E. casseliflavus/gallinarum* ASVs dominated in genotype WW and *E. mundtii* type in genotype yy (Fig. S10).

### Phylogenetic clustering and turnover of bacteria with life stage and genotype

We used measures of phylogenetic clustering and turnover to identify potential ecological mechanisms structuring the bacterial community with life stage and genotype. Bacterial communities in larval frass before and after transplantation and in adult abdominal fluid showed higher phylogenetic clustering than by chance (NRI, NTI > 0, Fig. 6A, 6B), which suggests selection or environmental filtering of specific bacterial lineages. In newly hatched larvae and adult gut, NRI and NTI did not differ from 0 and thus no evidence of selection of specific lineages was detected. Only frass before and after transplant showed potential differences in the extent of clustering with genotype. When looking at phylogenetic turnover in transitions between life stages, bacterial community turnover was largest in the transition from newly hatched larvae to frass (Fig. 6C). βNTI values of > 2 indicated deterministic selection in this community shift. Community dynamics of the other transitions appeared more driven by stochastic processes (−2 < βNTI > 2). Difference in βNTI values between the genotypes in frass suggests that genotype affected the dynamics and community assembly of the frass bacterial community.

**Fig. 6.**
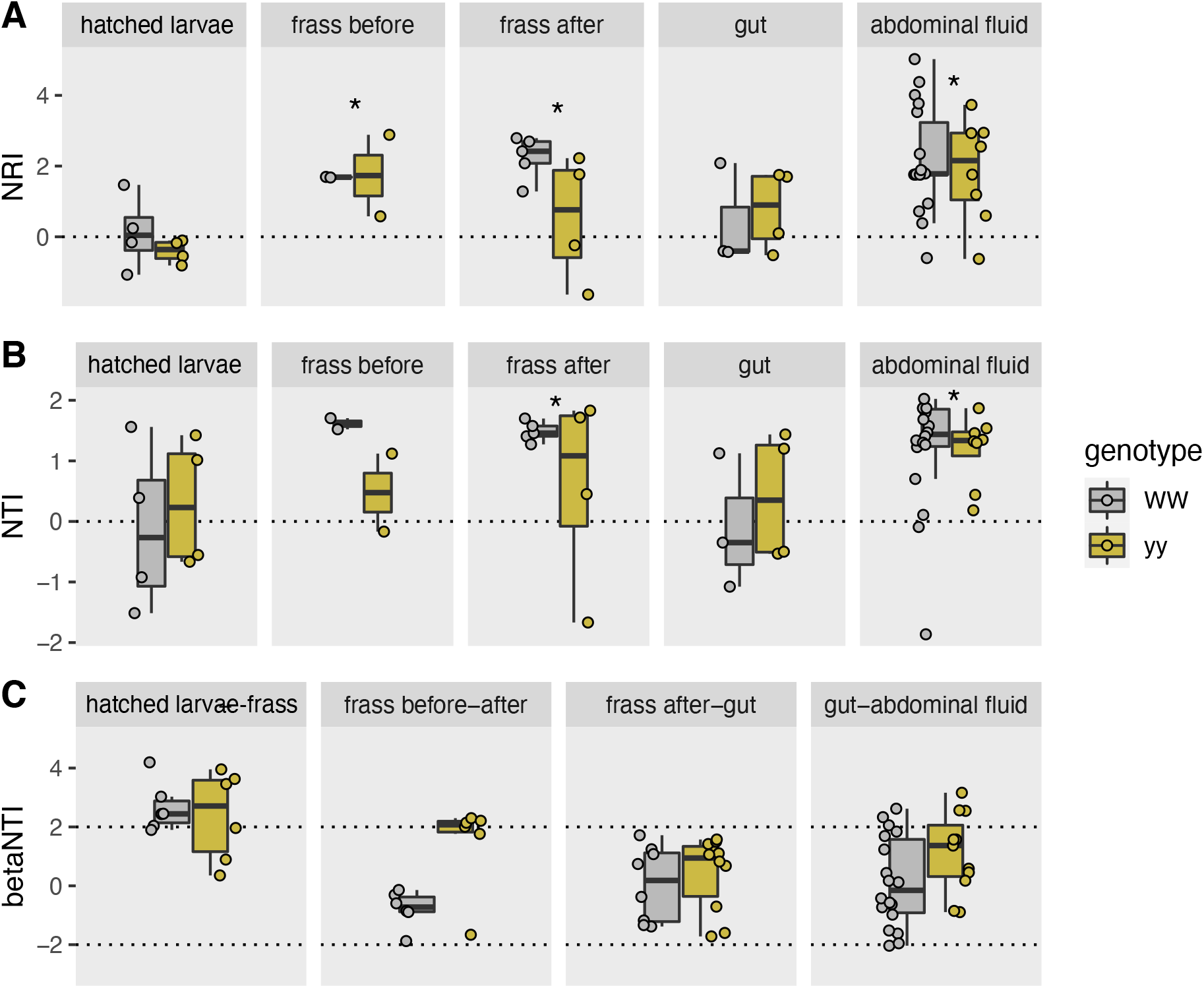
A) Net relatedness index (NRI) and B) nearest taxon index (NTI) with life stage and genotype (WW, yy) C) and beta nearest taxon index (βNTI) for bacterial community data between different stages. NRI > 0 and NTI > 0 (indicated in A and B by * when significant difference from 0 in t-test) show greater phylogenetic clustering than by chance. βNTI > 2 indicates significantly higher community turnover than by chance driven by deterministic selection, whereas −2 < βNTI > 2 suggests stochastic community assembly.

## Discussion

Bacteria in insect hosts have been connected to fitness and development (Feldhaar & Gross 2009, Gupta & Nair 2020) and even to driving changes in host allele frequencies (Rudman et al. 2019). Many insects show highly species-specific microbiome composition, but Lepidoptera are considered to deviate from this pattern of insect phylosymbiosis (Mallott & Amato 2021 but see Hammer et al. 2020). Consequently, the mechanisms of bacterial community assembly and the role of bacteria in key traits such as growth and defensive mechanisms of Lepidoptera remain unresolved (Hammer et al. 2017, Panigua Voirol et al. 2018). Here, we show that a reciprocal frass transplantation between a fast-growing and a slow-growing genotype of the wood tiger moth reversed the growth rate, but only in the fast-growing genotype, also increasing the volume of defensive secretions. The host bacterial community had components that 1) were genotype specific, 2) passed strong environmental filtering in the gut, and 3) were retained through life stages. Together, these findings suggest that whether the microbiome influences Lepidoptera growth may depend on genotype-specific ability of the host to gain and retain specific bacteria.

The life-history traits of the fast-growing genotype WW were more susceptible to frass transplantation than those of the slow-growing genotype yy. In addition to growth rate, the white and yellow colour morphs that the genotypes represent differ in their immune response mechanisms. The yellow morph that the genotype yy represents has more effective lytic activity in its haemolymph (Nokelainen et al. 2013). This stronger innate immune response against bacterial invaders could partly explain why the slow-growing genotype yy was more resistant to frass transplant than the WW genotype. The immune system has been proposed as one of the mechanisms driving microbiome differences between hosts with different genetic backgrounds (Franzenburg et al. 2013, Stagaman et al. 2017, Kohl 2020). Furthermore, the costs of this stronger immune defence in genotype yy could be reflected in slower growth (Boots & Begon 1993, Siva-Jothy et al. 2005, Ardia et al. 2012).

Effective defence against bacterial invaders in genotype yy could have prevented it from acquiring bacteria from the frass transplant or the environment that were potentially beneficial for growth in the fast-growing genotype WW. Alternatively, or in addition, the chemical or physiological environment in the gut of genotype WW may favour the establishment of different bacteria (Kohl 2020). We detected *Erysipelotrichaceae* only in the fast-growing genotype WW, and their relative abundance decreased together with growth rate after yy frass transplantation. The transplantation increased *Enterococcus* ASVs seen in yy frass, which could imply transfer of these bacteria to the WW genotype. We detected no transfer of bacteria from WW to yy. Presumably, WW-specific *Erysipelotrichaceae* (increasing growth), yy-originating *Enterococcus* (constraining growth) or the balance between these groups could be a bacterial component associated with larval growth.

*Erysipelotrichaceae*, which can be considered genotype-specific core taxa here, commonly occur in insect guts (Tegtmeir et al. 2016). For example, they occur in dung beetles depending on genus and diet (Ebert et al. 2021), and in Lepidoptera depending on growth environment (Gomes et al. 2020) and season (Higuita Palacio et al. 2021). The family *Erysipelotrichaceae* consists of anaerobic or aerotolerant bacteria with fermentative metabolism (Tegtmeier et al. 2016). However, the ASVs we detected are too distant from any described strains (Fig. S7) to allow more detailed speculation on their metabolism. In vertebrates, relative increase of *Erysipelotrichaceae* has been connected to dietary fat, weight gain, metabolic disorders, and high feed to weight conversion rate (Turnbaugh et al. 2008, Zhang et al. 2010, Kaakoush 2015, Stanley et al. 2016). Congruently, their decrease has been linked to reduced growth (Bao et al. 2020). These findings show *Erysipelotrichaceae* as a responsive member of the gut microbiota with connections to lipid metabolism and growth. Here, we linked *Erysipelotrichaceae* to lepidopteran larval growth rate when the genetic background allowed the establishment of these bacteria in the gut. Characterization of isolates or genomes of this distinct novel cluster of *Erysipelotrichaceae* is required to establish potential mechanisms for how these bacteria affect growth, directly or indirectly.

The other scenario that our results put forward is that specific members of *Enterococcus* could have an impact in reducing larval growth rate. Enterococci are lactic acid bacteria well adapted to survival in the harsh gut environment by evading host defences (Fiore et al. 2019, Mazumdar et al. 2021) and commonly found in Lepidoptera throughout their life cycle (Hammer et al. 2014, Chen et al. 2016, Teh et al. 2016, González-Serrano et al. 2020). The specific enterococci we detected in connection to slower growth clustered with *E. casseliflavus* and *E. gallinarum*, which can be the dominant bacteria in Lepidoptera (Martin & Mundt 1972, Tang et al. 2012, Shao et al. 2014, Jones et al. 2019). These enterococci have been suggested to potentially play beneficial roles for larvae, such as detoxifying diet plant compounds (Vilanova et al. 2016) and degrading cellulose and proteins (Pilon et al. 2013, Dantur et al. 2015), which may not align with reduced growth. On the other hand, strains of *E. casseliflavus* have also been reported to be pathogenic to larvae (Thakur et al. 2015, Shao et al. 2017), though not in all cases (Osborn et al. 2002). The larvae in our experiment showed no signs of pathogenic effects, and the only negative outcome we can connect the presence of *E. casseliflavus/gallinarum* is slower larval growth. In the control transplant of the faster growing WW genotype, we instead detected enterococci that grouped with *E. mundtii*. This proposed lepidopteran symbiont has been reported to be antagonistic against potential pathogens including *E. casseliflavus* by producing an antimicrobial peptide and to appear later in larval development than *E. casseliflavus* (Jarosz 1979, Johnston and Rolff 2015, Shao et al. 2017). Overall, our results suggest that the establishment and dynamics of these two clusters of enterococci (*E. casseliflavus/gallinarum* vs. *E. mundtii*) can be influenced by host genotype with potential effects on larval growth.

In adults of the wood tiger moth which, in contrast to larvae, are short-lived and do not feed, we detected both *E. mundtii* and *E. casseliflavus/gallinarum* ASVs across genotypes and treatments, which implies that different mechanisms could control their community dynamics in adults vs. larvae. If these dynamics are further sensitive to the genetic background of the host, as our results suggest, the specific functional roles of these enterococci are most likely highly context dependent. It is possible that under some conditions they form a host-adapted core microbiome (Shapira 2016, Risely 2020), which has often been considered to be lacking in Lepidoptera. The context-dependency would, however, suggest that any effects of these enterococci on growth are causal role functions depending on the presence or absence of the microbe, rather than selected effect functions driven by evolutionary mechanisms and consistent maintenance of the microbe in the host population (Klassen 2018). Further manipulative studies targeted at enriching or depleting such taxa are needed to confirm this notion.

Our results agree with previous findings on the low bacterial diversity in Lepidoptera and the lack of a consistent core microbiome with striking variability among individuals (Minard et al. 2019, Jones et al. 2019, Martínez-Solís et al. 2020). The artificial diet that we fed the larvae to minimize the presence of non-frass microbes most likely led to even lower diversity than in wild-collected or plant-fed larvae. This low bacterial load could also accentuate stochastic processes such as ecological drift and priority effects more than in hosts with more diverse and stable microbiomes, leading to high inter-individual variation (Chase and Myers 2011, Kohl 2020). Yet against this low and variable background, the results also support previous notions that specific taxa have an advantage in the harsh conditions of the lepidopteran larval gut (Tang et al. 2012). Taxa characteristic to a genotype (*Erysipelotrichaceae*) or common in Lepidoptera (*E. casseliflavus/gallinarum, E. mundtii*), which were not detected or barely detected in the most diverse bacteria of the newly hatched larvae, became dominant in larval frass via non-random selection and were retained in adults (Fig. 3, Fig. 6). Moreover, the filtering process in the larvae appeared to be largely genotype specific. In adults, the deterministic and genotype-specific patterns seen in larvae were attenuated, which may reflect the role of the adult stage in the life history of the wood tiger moth as a very short reproductive stage (5-7 days). A caveat here is that our set of adult gut samples is relatively small due to difficulties in dissecting them. Since adults do not feed, functional guts are not necessary, and gut structures are the remnants of the larval gut that are partly absorbed during metamorphosis. Nevertheless, our results provide a rare view on lepidopteran bacterial community dynamics through life stages in a species that does not feed as an adult. Butterflies that do feed as adults display relatively more consistent bacterial community composition in adults compared to larvae (Hammer et al. 2020). In the non-feeding wood tiger moth, bacteria in both whole larvae and adults varied a greatly, and passage through larval gut to frass was the most selective step.

The frass transplantation impacted the volume of abdominal secretions in the same way as the growth rate of the genotypes. The WW genotyped significantly increased its defensive secretions, whereas the expected opposite decrease in the yy was not observed (Fig. 2). A recent study found no differences in the volume of abdominal secretions of wild-caught or laboratory-reared white and yellow wood tiger moths (Lindstedt et al. 2020). Here, we observed larger volumes in the yy control genotype relative to the WW control genotype, and a general increase in both genotypes after frass transplantation (Fig. 2). The different diets may partly explain the discrepancies between studies. For instance, laboratory-reared moths were fed dandelion (*Taraxacum* sp.) in the previous study, whereas an artificial diet was used here. Likewise, the larval diet of wild-caught adults is unknown. Hence, it is difficult to draw comparisons between absolute volumes of defensive secretions. However, it can be inferred that diet is not a major contributor to the of defensive secretions since the frass transplantation increased the volume of defensive secretions in both genotypes which fed on the same artificial diet mixed with frass of the other genotype.

Differences in antipredator efficacy have also been reported, where the abdominal secretions of yellow moths are more deterrent against ants than secretions from white individuals (Rojas et al. 2019). The exact chemical composition of the abdominal fluids has not been characterised. However, the wood tiger moth can sequester hepatoxic organic compounds, such as pyrrolizidine alkaloids (PAs), from its diet (Winters et al. 2021), and these compounds have been detected in the abdominal secretions of wild-caught adults (Winters et al., unpublished). PAs are well-known for providing protection to plants against herbivores, which in turn can host bacterial associates with detoxifying abilities (Joosten et al. 2011, Tamariz et al. 2018). Some bacteria have even been suggested to synthesise PAs, as inferred by the presence of alkaloid biosynthesis gene clusters (Schimming et al. 2015). Manipulative experiments are needed to study PA sequestration-detoxification processes in the wood tiger moth. The great diversity of bacteria found in the abdominal secretions holds great potential to help elucidate the mechanism underlying these processes and to better understand differences in protection between the colour morphs.

The great and largely unexplained variation of bacteria among individuals in the defensive secretions represents an example of how the drivers of bacterial community composition can be decoupled in larvae vs. adults (Hammer and Moran 2019). We have previously shown that pre- and post-metamorphosis life stages in the wood tiger moth are only partly decoupled (Galarza et al. 2019). Our across-life stage analyses provide an additional line of evidence for this partial de-coupling. For instance, the genotype-specificity of *Erysipelotrichaceae* remained in the abdominal fluids, which showed that the bacterial community was not completely uncoupled from the previous life stages. A curious observation was a higher diversity of *Enterococcus* ASVs in the abdominal fluids compared to larvae, frass or gut. Together with the signs of phylogenetic clustering in the fluids, this finding suggests that distinct but partly unidentified drivers govern the bacterial community dynamics in the adult’s defensive secretions vs. in the larvae.

## Conclusions

We show for the first time that genotype-specific life-history traits in a lepidopteran host can be reversed with a reciprocal frass transplantation. Our results help clarify bacterial community assembly in Lepidoptera at different stages and relate the bacterial community composition to the genotype and growth of the wood tiger moth. With nearly full-length 16S rRNA gene amplicons, we were able to discern dynamics of closely related enterococci and recovered a novel cluster of insect-associated, genotype-specific *Erysipelotrichaceae*.

Our findings indicate that an insect host can fine-tune its bacterial community in a genotype-specific manner even against a background of a highly variable bacterial community with stochastic community assembly. The strong selection of bacteria in larval frass appeared to be relaxed in the adults, which shows that previous findings of consistent bacterial community in adult Lepidoptera (Hammer et al. 2020) may not apply to species that do not feed as adults. Thus, not only life stage but also species-specific adult feeding habits and host genotype can influence the bacterial community assembly contributing to the high variation and inconsistencies observed in Lepidoptera microbiomes. Overall, our results suggest that the digestive processes of slow- and fast-growing host genotypes differ in filtering or retaining specific bacterial groups. Looking deeper into the functional, chemical and genomic background of these differences could reveal the molecular and ecological mechanism of the filtering.

## Declarations

### Ethics approval and consent to participate

Not applicable

### Consent for publication

Not applicable

### Availability of data and material

The sequence data are available in the National Center for Biotechnology Information Sequence Read Archive under accession code PRJNA804133. R scripts and data files for microbiome analysis are available at https://github.com/helijuottonen/mothtransplant. R scripts and data files for life history analysis will be available at https://github.com/Juan-Galarza?tab=repositories.

### Competing interests

The authors declare that they have no competing interests.

### Funding

This project was funded by the Academy of Finland grant 322536 to JAG and 328474 and 345091 to JM.

### Authors’ contributions

JAG, JM and NNM design the study; JAG, NNM and LM carried out the experiments and laboratory work; JAG analysed life history data; HJ analysed microbiome data; HJ, JAG and NNM wrote the manuscript. All authors commented and approved the manuscript.

## Acknowledgements

We thank Sari Viinikainen and Elisa Salmivirta for assistance in laboratory work and the Finnish IT Center for Science for computational resources.

## Notes

### Competing Interest Statement

The authors have declared no competing interest.

