## Supplementary materials for "Host’s genetic background determines the outcome of reciprocal faecal transplantation on life-history traits and microbiome composition"

§ equal contribution
* corresponding author

**Contents:**

Supplementary figures:

Fig. S1 Larval developmental time in laboratory stocks of the genotypes
Fig. S2 Normality and heterogeneity of life history traits
Fig. S3 Developmental time of *Arctia plantaginis* in reciprocal frass transplant
Fig. S4 Pupal weight of *A. plantaginis* with reciprocal frass transplant
Fig. S5 Phylogenetic diversity ASV number with tissue type and genotype
Fig. S6 NMDS of bacterial communities based on MPD
Fig. S7 Phylogenetic tree of *Erysipelotrichaceae* ASVs
Fig. S8 Phylogenetic tree of selected *Enterococcus* ASVs
Fig. S9 Taxonomic composition of bacteria in diet samples
Fig. S10 Taxonomic composition of bacteria in newly hatched larvae, gut and abdominal fluid

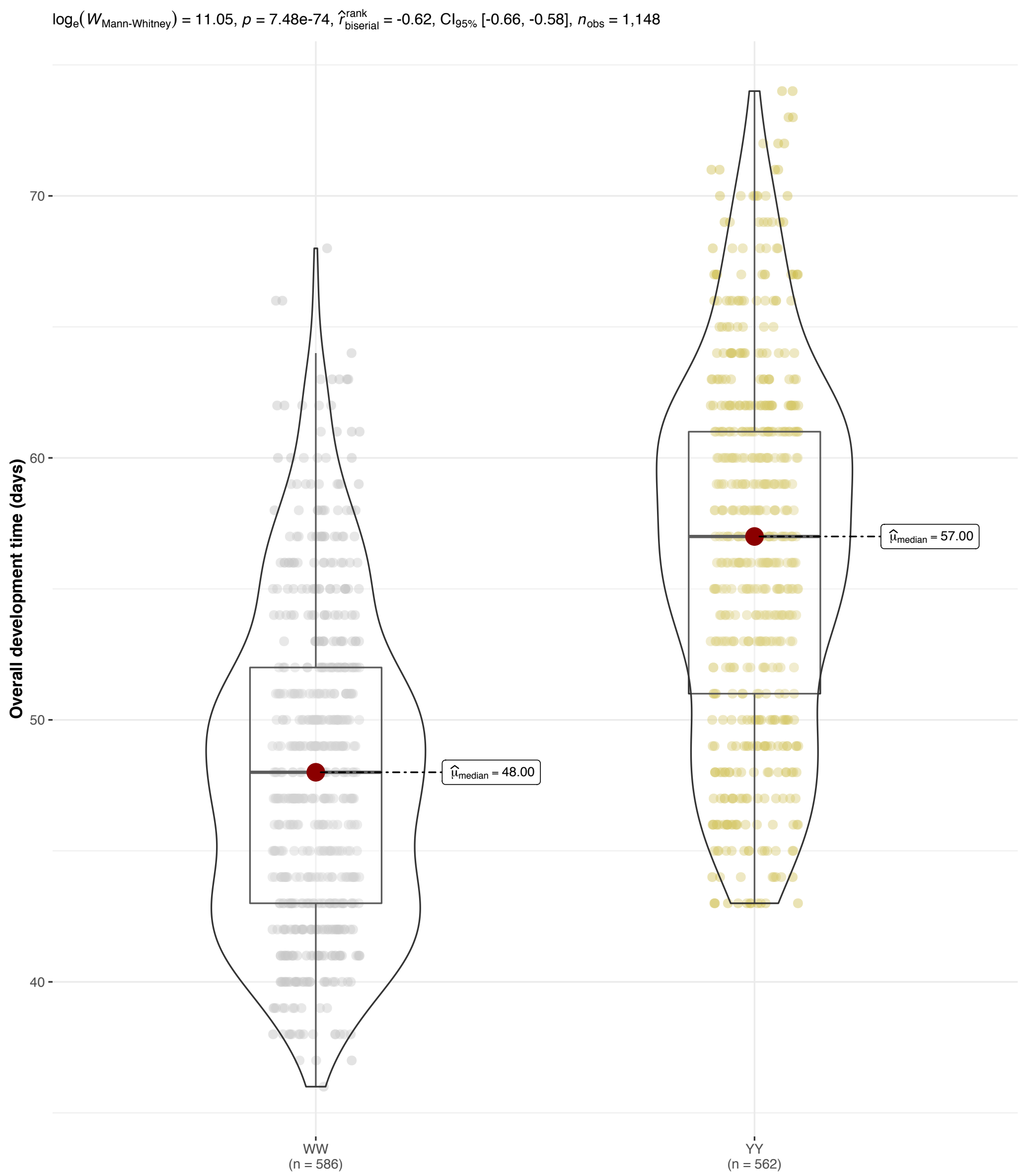


**Fig. S1.** Larval developmental time of wood tiger moth (*Arctia plantaginis*) genotypes reared in laboratory conditions. Shown are the results of one-way ANOVA test of equal means and Student's *t*-test with P-values adjusted for multiple comparisons.

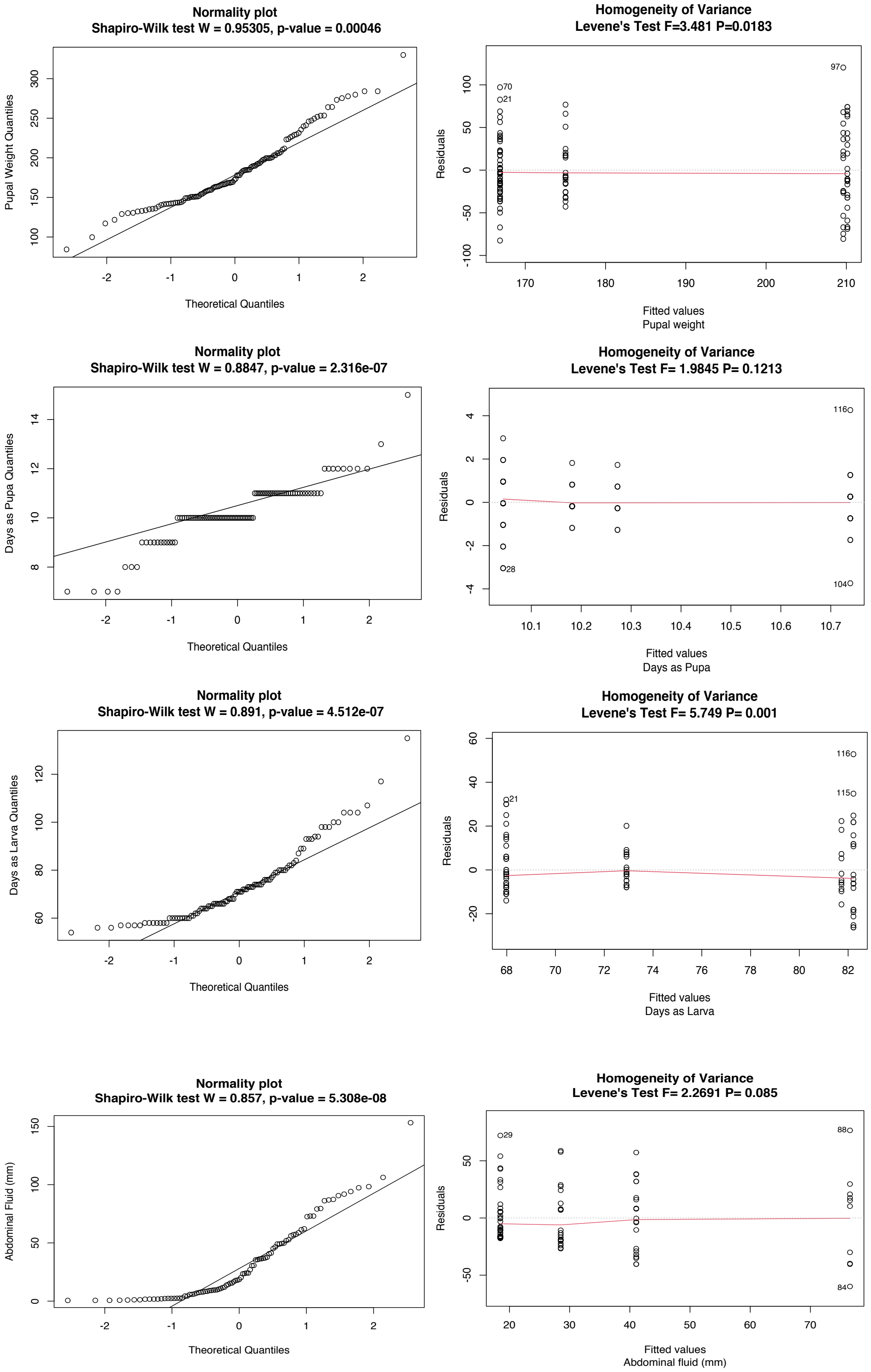


**Fig. S2**. Normality and homogeneity of variance in wood tiger moth (*Arctia plantaginis*) life-history traits. Shown are the results of Shapiro-Wilk test for normality and Levene's test for homogeneity of variances.

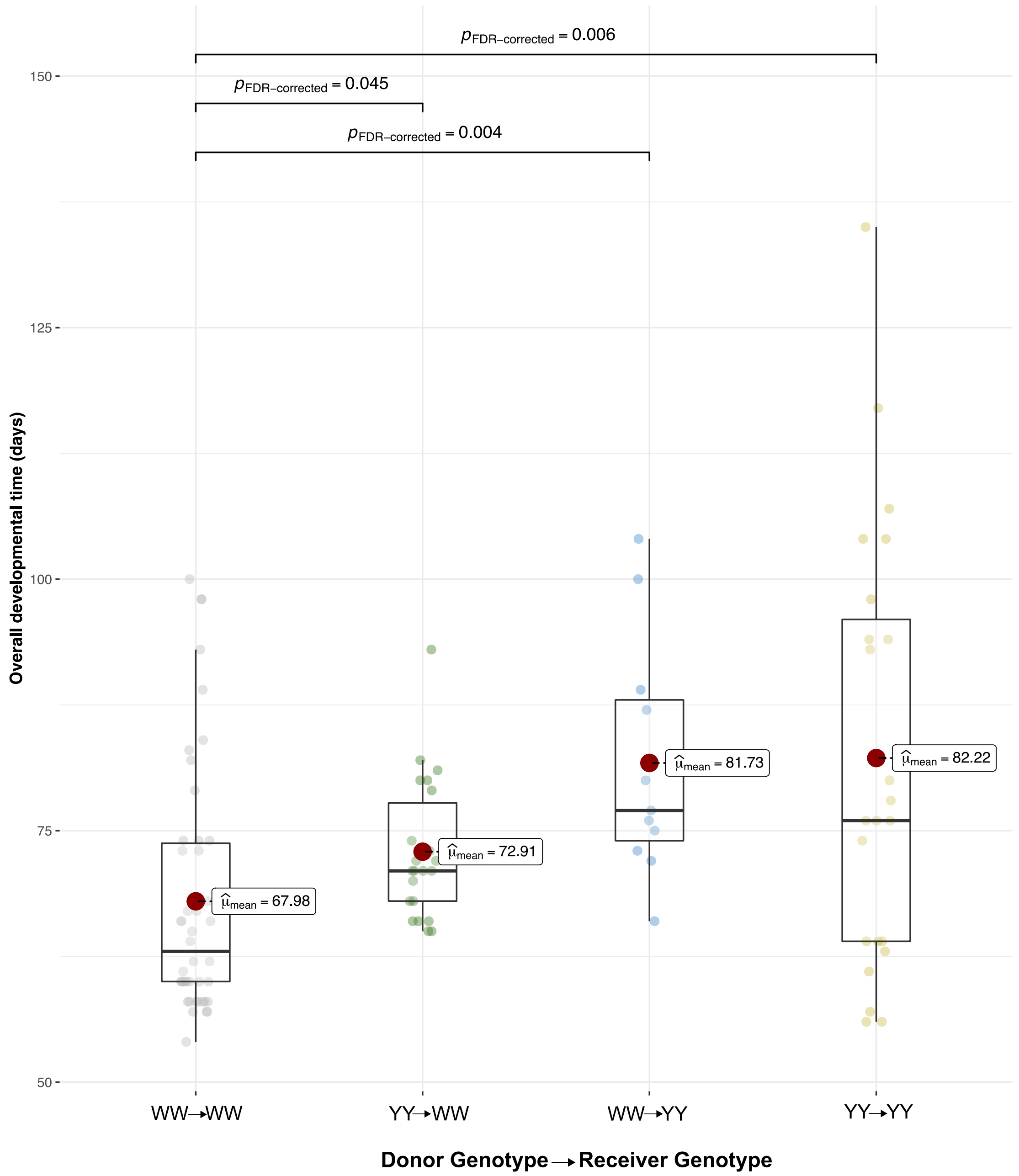


**Fig. S3.** Developmental time of *A. plantaginis* in reciprocal frass transplant between genotypes WW and yy and as controls within the genotypes.

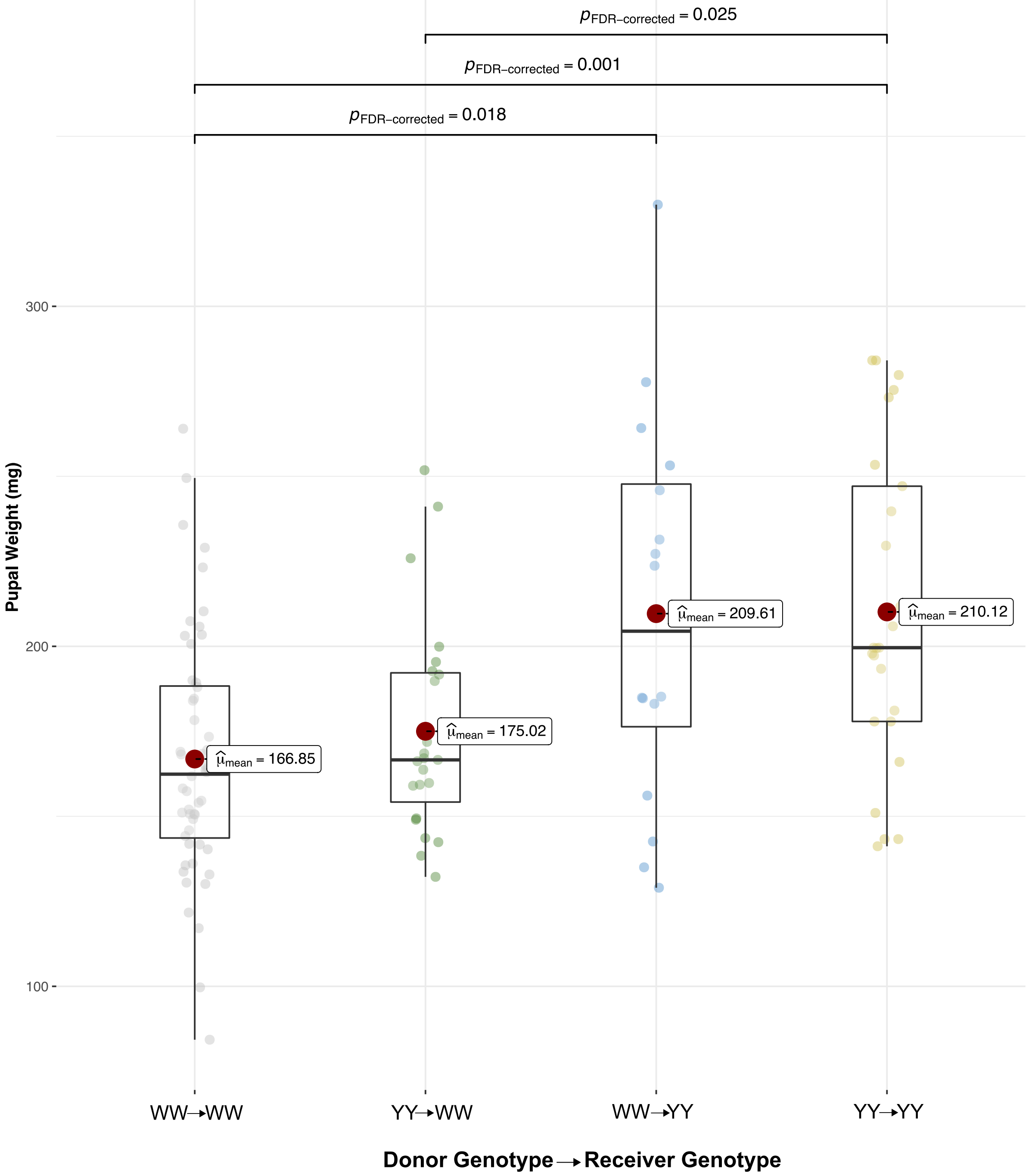


**Fig. S4.** Pupal weight of *A. plantaginis* with reciprocal frass transplant between genotypes WW and yy and as controls within the genotypes.

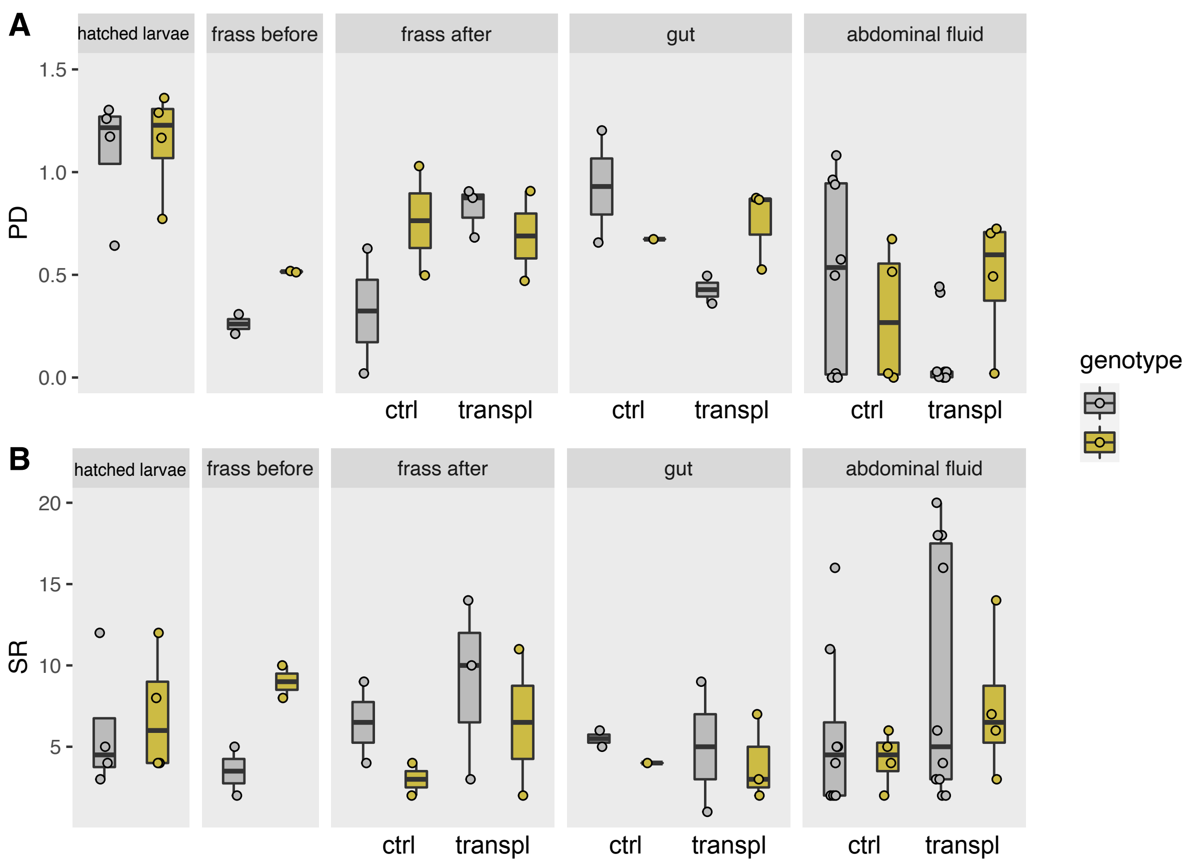


**Fig. S5** A) Phylogenetic diversity (PD) and B) ASV number (SR) with tissue type and genotype. For tissues collected after frass transplantation, diversity is shown in control transplantations within genotypes (ctrl) and transplantations between genotypes (transpl).

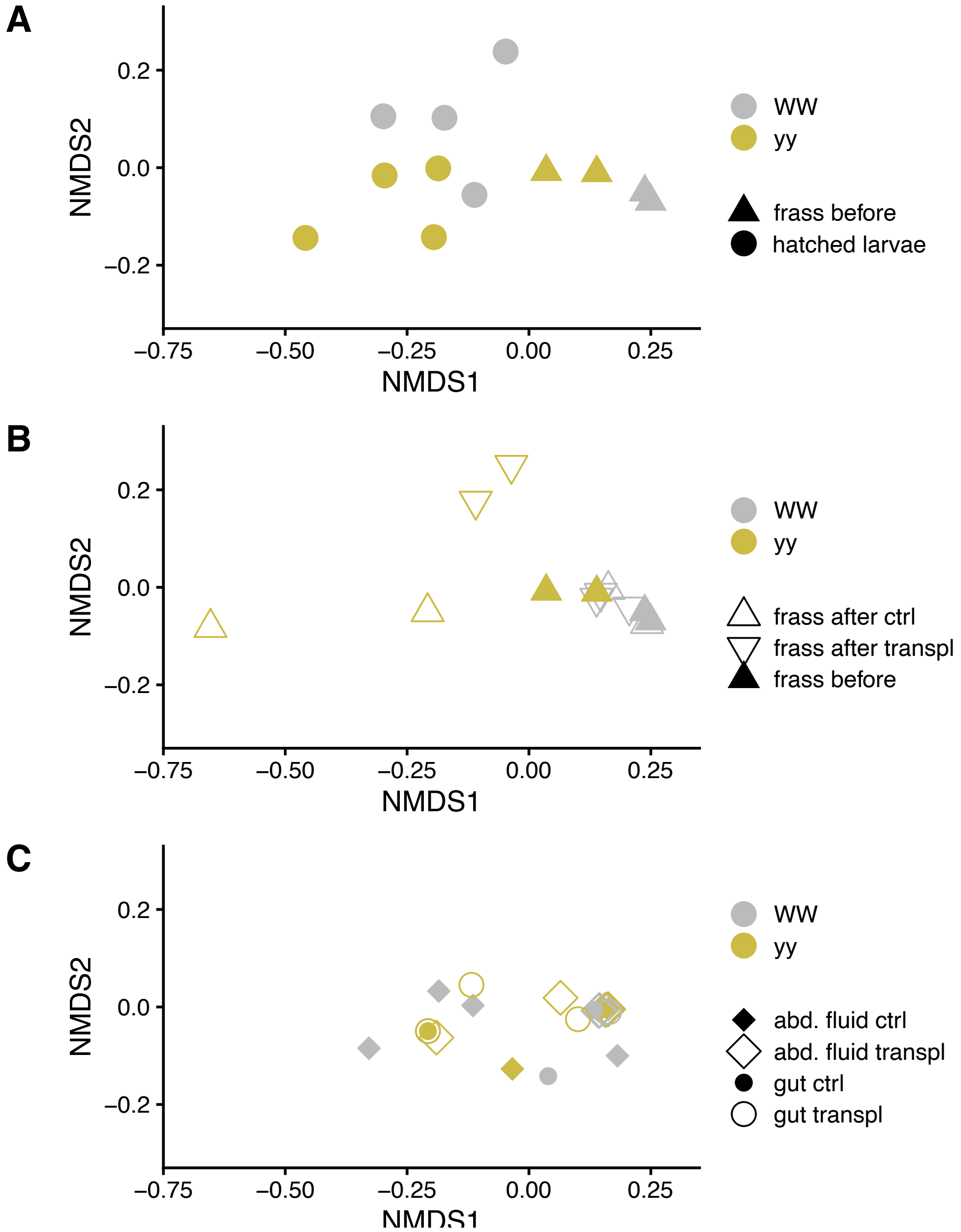


**Fig. S6.** Non-metric multidimensional scaling (NMDS) plots of bacterial community based on phylogenetic distances (MPD, mean pairwise distance) and separated by moth genotype (WW, yy). A) Whole larva and frass before transplantation, B) frass before and after transplantation (control transplantation within genotypes and transplantation between genotypes WW and yy), C) gut and abdominal fluid. Panels A, B and C are based on the same ordination but plotted separately. Stress = 0.07.

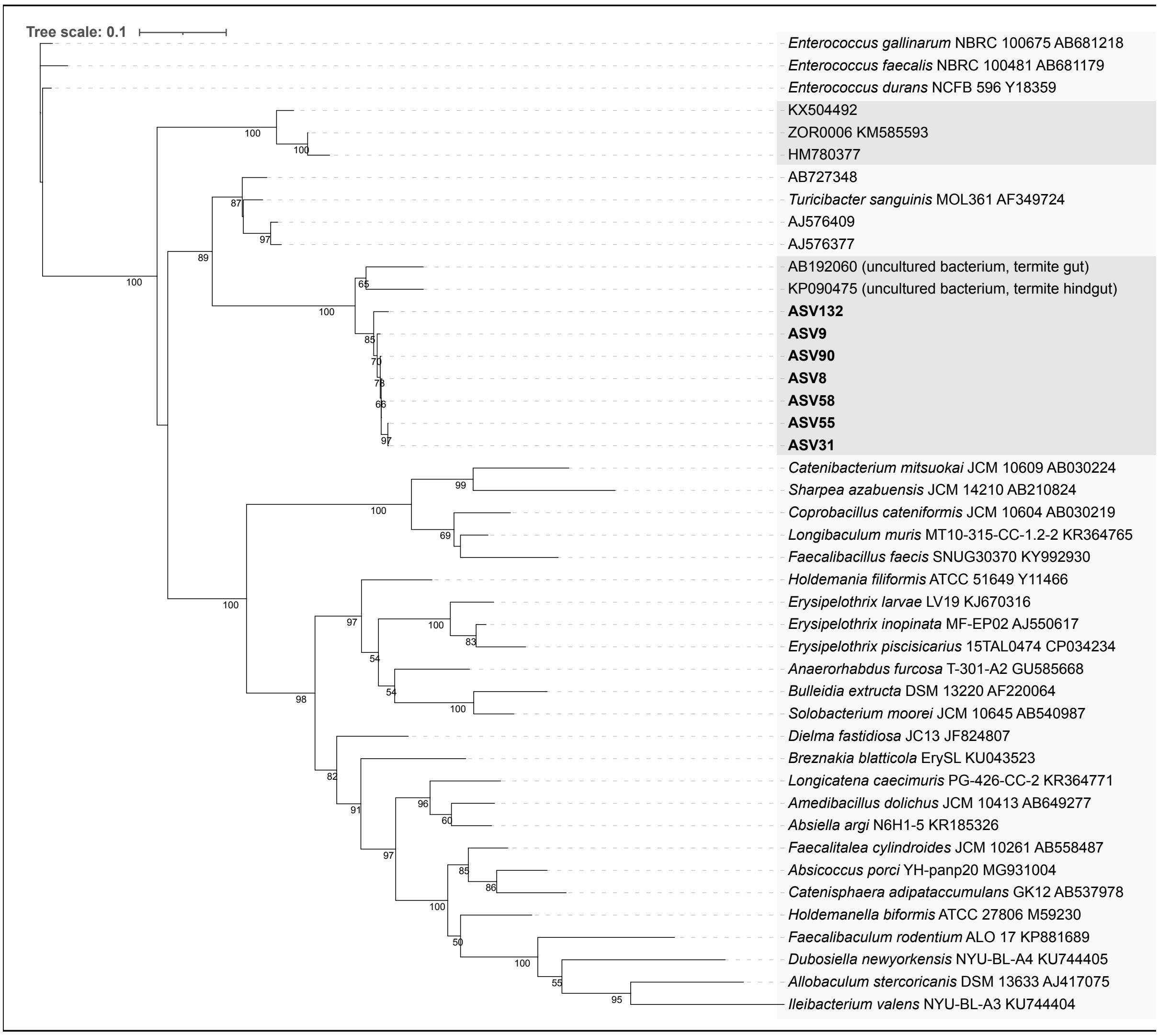


**Fig. S7.** Maximum likelihood tree of *Erysipelotrichaceae* ASVs (in **bold**) and selected reference sequences based on nearly full length 16S rRNA gene sequences (alignment length 1415 bp). Bootstrap values were determined with 100 replicates in RAxML. Scale bar indicates 10% sequence divergence.

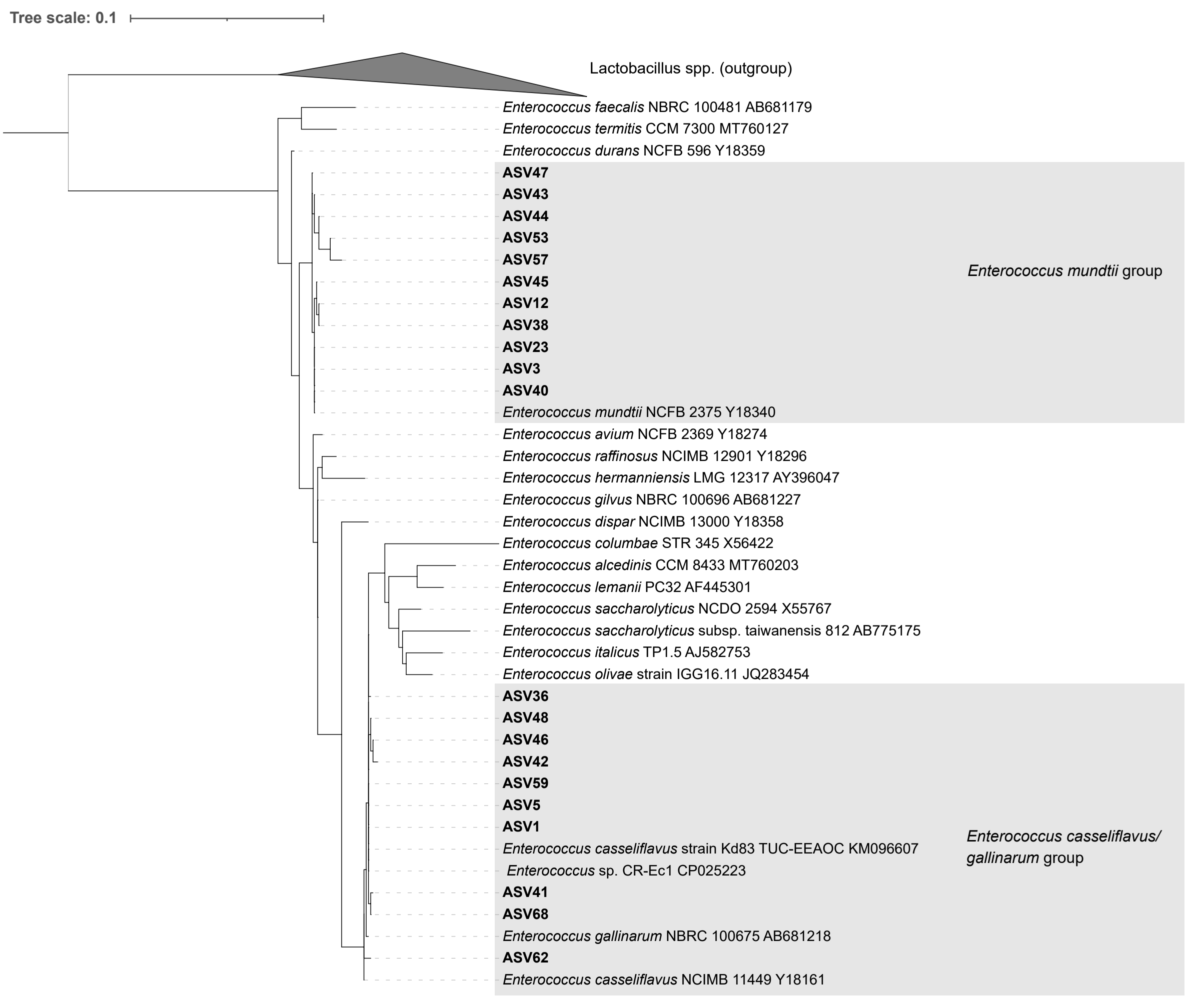


**Fig. S8.** Maximum likelihood tree of *Enterococcus* ASVs (in **bold**) and selected reference sequences (*Enterococcus* type strains and additional strains or genomes with the highest similarity in Blast to major ASVs) based on nearly full length 16S rRNA gene sequences (alignment length 1270 bp). Bootstrap values were determined with 100 replicates in RAxML. Scale bar indicates 10% sequence divergence.

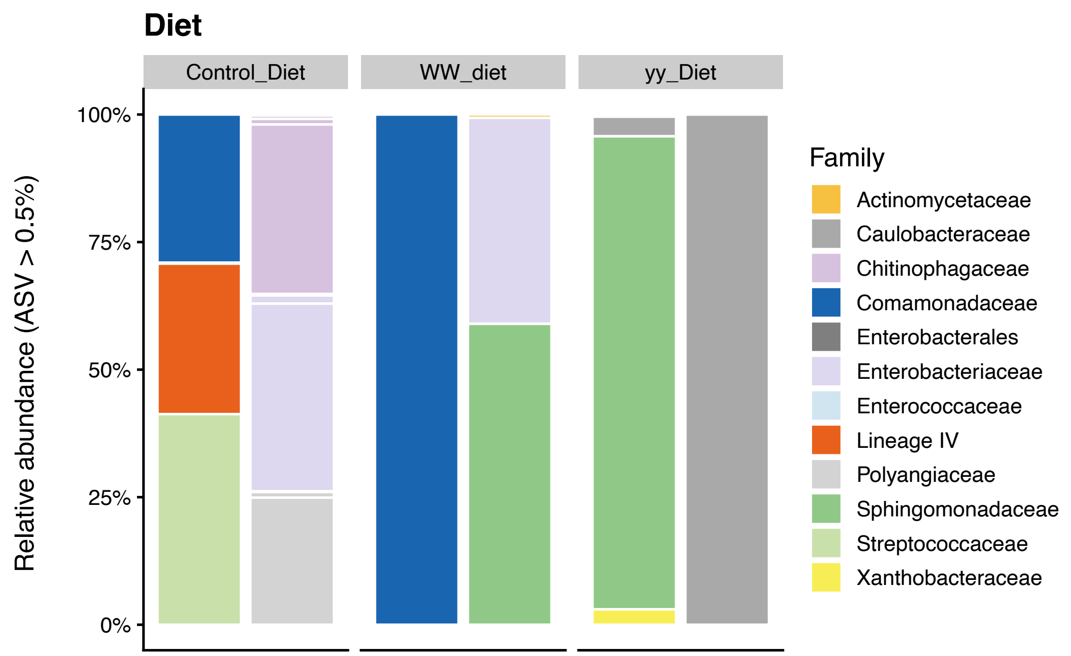


**Fig. S9**. Taxonomic distribution of bacterial 16S rRNA gene amplicon sequence variants (ASVs) at family level in control diet and in transplant diets. Each bar represents one sample. White horizontal lines separate ASVs and each ASV with relative abundance above 0.5% is shown as its own section in the columns. WW diet stands for control diet mixed with frass of genotype WW. Yy diet stands for control diet mixed with frass of genotype yy). Lineage IV belongs to phylum *Elusimicrobiota*.

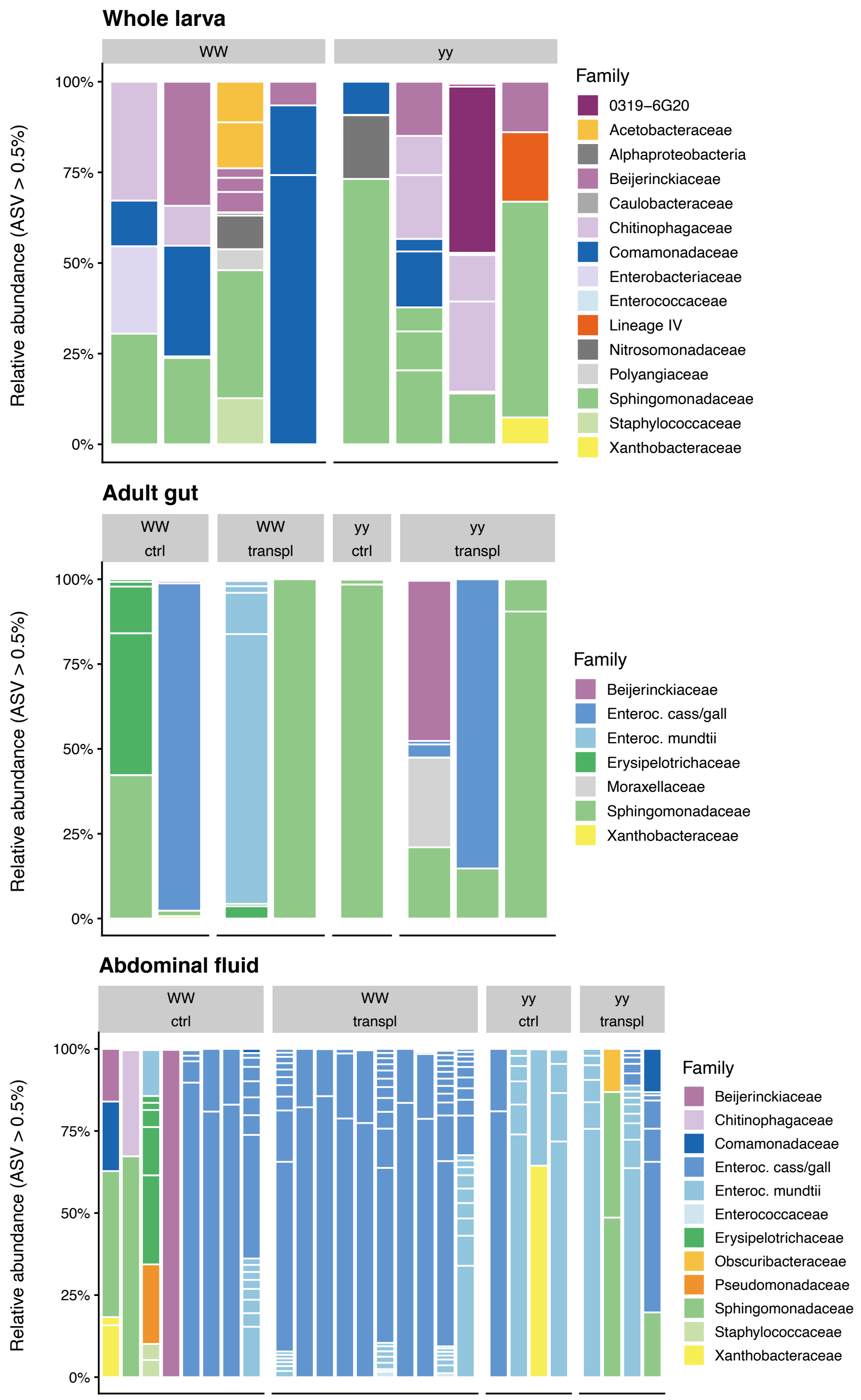


**Fig. S10.** Taxonomic distribution of bacterial 16S rRNA gene amplicon sequence variants (ASVs) at family level in surface-sterilized hatched larvae, adult gut and adult abdominal fluids in *Arctia plantaginis* genotypes WW and yy. Each bar represents one sample. White horizontal lines separate ASVs and each ASV with relative abundance above 0.5% is shown as its own section in the columns. Ctrl, control transplants that received the frass of their own genotype. Transpl, transplants that received frass of the other genotype. Lineage IV belongs to phylum *Elusimicrobiota* and '0319-6G20' to phylum *Bdellovibrionota*, class *Oligoflexia*. Enteroc. stands for *Enterococcus*, and cass/gall for *casseliflavus/gallinarum*.
